# The Molecular Basis of Specific DNA Binding by the BRG1 AT-hook and Bromodomain

**DOI:** 10.1101/854000

**Authors:** Julio C. Sanchez, Liyang Zhang, Stefania Evoli, Nicholas J. Schnicker, Maria Nunez-Hernandez, Liping Yu, Jeff Wereszczynski, Miles A. Pufall, Catherine A. Musselman

## Abstract

The ATP-dependent BAF chromatin remodeling complex plays a critical role in gene regulation by modulating chromatin architecture, and is frequently mutated in cancer. Indeed, subunits of the BAF complex are found to be mutated in >20% of human tumors. The mechanism by which BAF properly navigates chromatin is not fully understood, but is thought to involve a multivalent network of histone and DNA contacts. We previously identified a composite domain in the BRG1 ATPase subunit that is capable of associating with both histones and DNA in a multivalent manner. Mapping the DNA binding pocket revealed that it contains several cancer mutations. Here, we utilize SELEX-seq to identify the DNA specificity of this composite domain and NMR spectroscopy and molecular modelling to determine the structural basis of DNA binding. Finally, we demonstrate that cancer mutations in this domain alter the mode of DNA association.

## Introduction

The eukaryotic genome exists in the nucleus in the form of chromatin. At its most basic level chromatin is composed of iterative subunits known as nucleosomes [1], which consist of an octamer of histones (two each of H2A, H2B, H3, and H4), wrapped by ∼147 base pairs of DNA. Chromatin structure needs to be spatially and temporally remodeled for all DNA templated processes. This is in large part aided by ATP-dependent remodeling complexes. These can be categorized into four families and include the SWI/SNF, ISWI, IN080, and CHD families of remodelers. The BRG1/BRM Associated Factors complex (BAF complex) is the human homologue of yeast SWI/SNF. BAF is highly polymorphic, forming several different canonical and non-canonical complexes in embryonic stem cells as well as upon differentiation [2–4]. Additional heterogeneity is seen depending on developmental stage and cell type. BAF is important for sliding and ejecting histones in a variety of cellular processes, including transcription regulation and DNA repair [5].

BAF complexes are tailored for regulation at specific chromatin regions. The exact mechanism by which BAF is recruited to these regions is not fully understood. It is in part mediated by association with transcription factors. In addition to this, its many subunits harbor a number of histone and DNA binding domains [6, 7]. These domains are thought to form a large multivalent network of chromatin contacts, that guide its association with the proper nucleosome substrates, and regulate remodeling activity. These include readout of histone post-translational modifications as well as specific and non-specific recognition of DNA elements. However, the details of these interactions are still being uncovered, as is the mechanism by which they coordinate to promote proper recruitment and function.

BAF subunits are mutated in greater than 20% of all human cancers sequenced to date, making it one of the most frequently mutated chromatin regulatory complexes. Mutations are found throughout most of the subunits, but are especially enriched in BAF250A and BRG1 (see below) [7]. While most mutations lead to loss of function of that subunit, the mutated complexes are potentially good therapeutic targets. Thus, an understanding of the mechanism by which BAF associates with chromatin is critical both for a fundamental understanding of the complex as well as for determining etiology of disease and in the development of pharmaceutic interventions.

The BAF complex contains one of two possible catalytic subunits, either BRG1 or BRM. These are mutually exclusive ATPases in the BAF complex. Although highly homologous, they cannot substitute for one another *in vivo* [8]. This is in part due to association with distinct transcription factors through their unique N-terminal domains[9]. BRG1-containing BAF is critical for development, with mutation of the ATP binding site in BRG1 being embryonic lethal in mice [10]. Furthermore deletion of BRG1 in mouse intestine leads to early death [11].

Both BRG1 and BRM contain a highly conserved bromodomain (BD) at the C-terminus (Figure 1A), the presence of which is unique among remodeler ATPases. Both bromodomains (BDs) have previously been characterized to bind acetylated-lysine on histone tails, with a moderate preference for histone H3 acetylated at lysine 14 (H3K14ac)[12–14]. In addition to its histone binding activity, we have recently discovered that both BDs can associate with DNA [15]. Nucleic acid binding activity has only been demonstrated for a few BDs [15, 16]. However, we have predicted that ∼30% of known BDs will bind to nucleic acids [15], highlighting the importance of better understanding this function. Previous studies indicate that the BRG1 BD is not essential for global chromatin association of the BAF complex. For example, small molecule inhibitors that target the BRG1 BD acetyl-lysine binding pocket failed to disengage the ATPase from chromatin, except in the case of hyperacetylation [17]. Similarly, mutation of the DNA binding interface does not alter the affinity of BRG1 for bulk chromatin purified out of mouse embryonic stem cells (ESCs). However, ESCs treated with the BRG1 BD inhibitor, PFI-3, showed a reduced stemness potential [18]. In addition, both the acetyl-lysine and DNA binding pockets are conserved and mutated in cancer. Together, this suggests that the BD is critical for BAF function, which could include targeting or retention to specific binding sites, and/or regulation of ATPase or remodeling activity.

**Figure 1.**
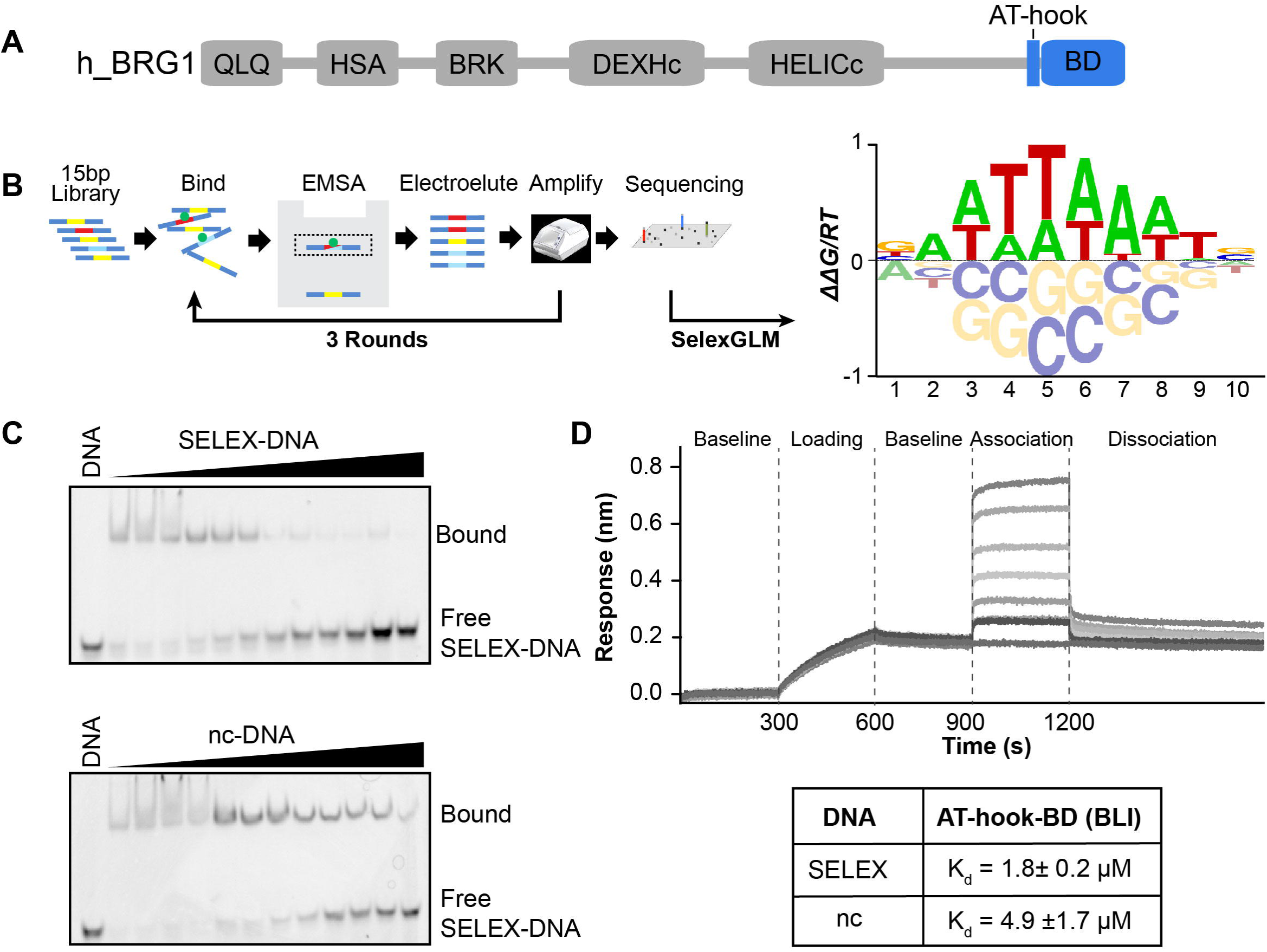
DNA specificity of the BRG1 ATBD. (A) Domain architecture of the human BRG1 subunit of the BAF complex. The AT-hook and bromodomain (BD) are highlighted in blue. (B) Schematic representation of SELEX-seq followed by a construction of a position specific affinity matrix and converted into the affinity logo containing AATTAAAT. (C) Specificity was confirmed through competitive binding assays analyzed by EMSA. A 15bp double-stranded DNA containing either the SELEX determined DNA sequence (SELEX-DNA: CCTCAATTAAATCTC) or a non-consensus sequence (nc-DNA: CCTCAGTCGGTCGTA) was used. The complex of ATBD and Cy5-labeled SELEX-DNA (bound) was competed against unlabeled SELEX-DNA (top) or nc-DNA (bottom). (D) Dissociation (K_d_) values were determined by BLI. Shown is a set of representative traces in a BLI binding assay (top). Calculated K_d_ values are in the table below, values reported are the average of 3 runs and standard deviations.

Ten residues N-terminal to the BD in both BRG1 and BRM is an AT-hook (Figure 1A). AT-hooks, described for the first time in the context of the high mobility group (HMG) proteins [19], are short motifs of between nine and ten amino acids that are arginine/lysine rich. A central consensus sequence of glycine-arginine-proline (GRP) allows the motif to insert into the minor groove of DNA. In addition, the extended AT-hook has been shown to bind to RNA [20]. AT-hooks prefer AT-rich DNA sequences as this leads to narrowing of the minor groove and a more focused electrostatic potential [21, 22]. In a genome wide study by Aravind et al. [23], many examples were found of an AT-hook existing just adjacent to another DNA-binding module, and it was suggested that this is a common auxiliary motif cooperating with adjacent structured DNA binding domains [23]. Similar to the BD, the role of the AT-hook in BAF function has not been elucidated. However, it is also highly conserved, and known to be mutated in cancer. We have recently shown that the tandem AT-hook and BD (AT-BD) function as a composite DNA binding module within the BAF complex, associating with DNA in a multivalent manner, with low micromolar affinity [15]. We demonstrated that the AT-BD can bind both naked and nucleosomal DNA. However, many questions remain including whether this module has sequence specificity, and the molecular basis of binding to DNA. Here, we utilize SELEX-seq to determine the sequence specificity of the BRG1 AT-BD and NMR spectroscopy and molecular modelling to uncover the structural basis of DNA binding.

## Materials and Methods

### Protein Expression and Purification

Human BRG1 AT-BD (residues 1434–1569) or BD (residues 1454-1569) were cloned into a pGEX6p-1 vector (GE Healthcare) from full length BRG1 cDNA obtained from GE Open Biosytems. Proteins were expressed in BL21 (DE3) *E. coli* cells (New England Biolabs, Ipswich, MA) in LB or M9 minimal media supplemented with vitamins (Centrum), 1 g/L ^15^NH_4_Cl and 1g/L ^12^C or ^13^C D-glucose. When the culture reached OD_600_=1, protein expression was induced with 0.16 mM isopropyl β-D-1-thiogalactopyranoside (IPTG) for 16 hours at 28^⁰^C. Cells were lysed with a homogenizer (Avestin Emulsiflex C3) in resuspension lysis buffer containing 20 mM Tris pH 7.5, 500 mM KCl, 2.5 mM MgSO_4_, 0.5% triton, 3mM DTT, 1mg/ml lysozyme, DNaseI, and a protease inhibitor cocktail tablet (Roche). The lysate was cleared through centrifugation for 45 min at 12,000*×*g. The resulting supernatant was incubated with glutathione agarose resin (ThermoFisher Scientific) for one hour. The resin was washed thoroughly with buffer containing 50 mM Potassium phosphate pH 7.0, 50 mM KCl, and 2 mM DTT (final buffer). The GST-tag was cleaved off with PreScission Protease, and the cleaved protein was further purified using FPLC first with cation exchange (Source-S, GE Healthcare), followed by gel filtration (Superdex 75, GE Healthcare).

### Selective Evolution of Ligand by Exponential Enrichment (SELEX)

Oligos used:

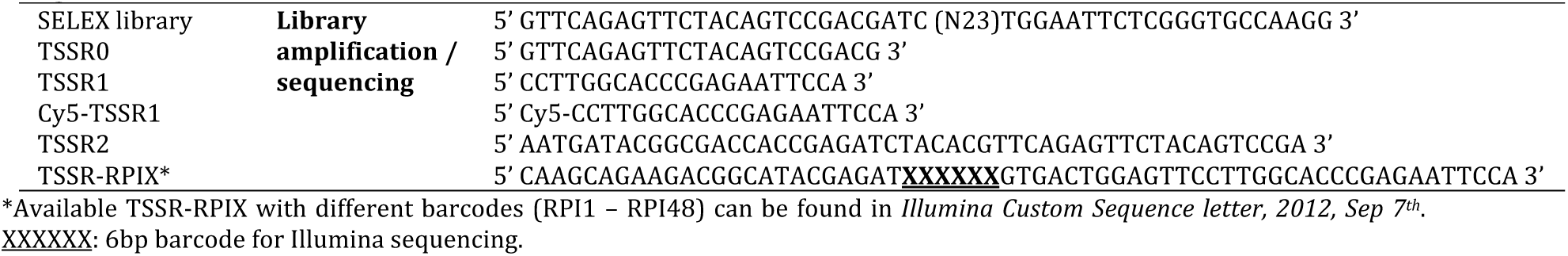

SELEX libraries consist of a 15-base pair (bp) randomized region flanked by TruSeq Small RNA adapter sequences. The 62 bp libraries were ordered as a single strand in the 1 µM desalted format from Integrated DNA Technologies using the hand mixing option for the randomized region to ensure maximal randomization. The second strand was synthesized by Klenow extension using the Cy5-TSSR1. A 100µl reaction containing 2µM library, 5 µM Cy5-TSSR1, 150 µM dNTPs in New England Biolabs (NEB) buffer 2 was incubated at 94°C for 3 minutes, then cooled to 37°C over 45 minutes. After addition of 24 units of Klenow fragment, the reactions were incubated at 37°C for 60 minutes, 72°C for 20 minutes, and then cooled to 10°C over 45 minutes. The dsDNA library was purified using a MinElute column (Qiagen) using the standard protocol, and then quantified by absorbance at 260 nm.

SELEX-seq was then performed as previously described [24], but with a few modifications. For SELEX, the bound fraction of library was separated from free by electrophoretic mobility shift assay (EMSA). In a 125µl reaction, 0.5 µM AT-BD was incubated with 2µM library in binding buffer (20 mM TrisHCl, pH 8.0, 150 mM KCl, 5% Glycerol, 1 mM EDTA, 5 mM MgCl_2_, 40 ng/µl Poly(dIdC) (Sigma: P4925), 200 ng/µl BSA, 1 mM DTT, 200 mg/ml Ficoll PM400 (Sigma: F4375)) at 4°C for 1 hour. Reactions were then run on a non-denaturing PAGE (10% 75:1 acrylamide:bis-acrylamide, 0.2X TBE) for 50 minutes in 0.2X TBE at 125 V. Free and bound bands were visualized by Cy5 fluorescence (GE LAS4000 imager). The shifted bands were excised, and the DNA recovered by electroelution (Novagen D-tube dialyzer, 3.5 kDa) in 0.2X TBE. The recovered library was purified (Qiagen MinElute PCR clean-up) and eluted (10 mM TrisHCl, pH 8.0) in a final volume of 180µl. In order to maximally amplify the library while minimizing artifacts, recovered library was quantified by qPCR, divided into 85 reaction of 100 µl, and amplified using Phusion (NEB). The number cycles depended on how much DNA was present and was calculated as the number of cycles for 90% signal in qPCR plus one additional cycle. Three rounds of selection were performed in total.

Adapters with unique barcodes were added to the libraries barcodes by amplifying input (Round 0 or R0) and Round 3 (R3) using TSSR2 and TSSR-RPIX. For each library, 400ng of library was amplified for 2 rounds using standard Phusion (NEB) PCR conditions. The 133bp final library was then separated from the 63 bp input on a 16% native PAGE (0.5X TBE, 19:1 acrylamide/bis-acrylamide), excised, and recovered by electroelution. Libraries were then checked for quality and quantified by Bioanalyzed (Agilent Technologies), then sequenced to a depth of ∼10 million reads/library using single-end 50bp sequencing.

The sequenced libraries were initially processed using the *R* package *SELEX* (R Core Team)(http://bioconductor.org/packages/SELEX). To correct for biased sampling from the input library, a markov model was first determined (6^th^ order), then constructed for R0 using the selex.mm(). Next, the optimal motif length was defined by determining the Kullback-Leibler divergence for motifs of increasing size using selex.infogain(). The optimal motif length proved to be 10bp.

Using the R package SelexGLM (https://www.bioconductor.org/), an affinity table was first constructed for k=10 using selex.affinities (). An initial position specific affinity matrix (PSAM) was constructed (BAWTTWAWTS) and used as a seed. The subsequent iterative procedure alternated between two steps. First, the current PSAM was used to find the position/direction of highest affinity on either strand, the optimal “view” on the probe. If that optimal affinity was larger than 95% of the sum over all positions (including the top position), the probe was used in the analysis; otherwise, it was ignored. The set of optimal positions in each of the accepted probes was used to define a design matrix containing the base identity at each position relative to the start of the optimal view. Using the probe counts as independent variables, the logarithm of the expected probe frequency in R0 according to the Markov model as offset, and a logarithmic link function, a fit was performed using the glm () function. The regression coefficients were interpreted as free-energy differences ΔΔG. All computational figure panels in this paper were produced fully automatically from the raw sequencing data using *R* scripts that use the *SELEX* and *SelexGLM* packages.

### SELEX-DNA and nc-DNA Preparation and Purification

Oligonucleotides for BLI, NMR, and gel shift experiments were purchased from Integrated DNA Technologies (IDT) (see Table 1). Forward and reverse strands were resuspended in 50 mM potassium phosphate pH 7.0, 50 mM KCl, and 2 mM EDTA, in nuclease free water, mixed at equimolar ratios and annealed by heating to 94 ^⁰^C for 10 min before being slowly cooled to room temperature. The annealed strands were then further purified by gel filtration chromatography (Superdex 75, GE Healthcare).

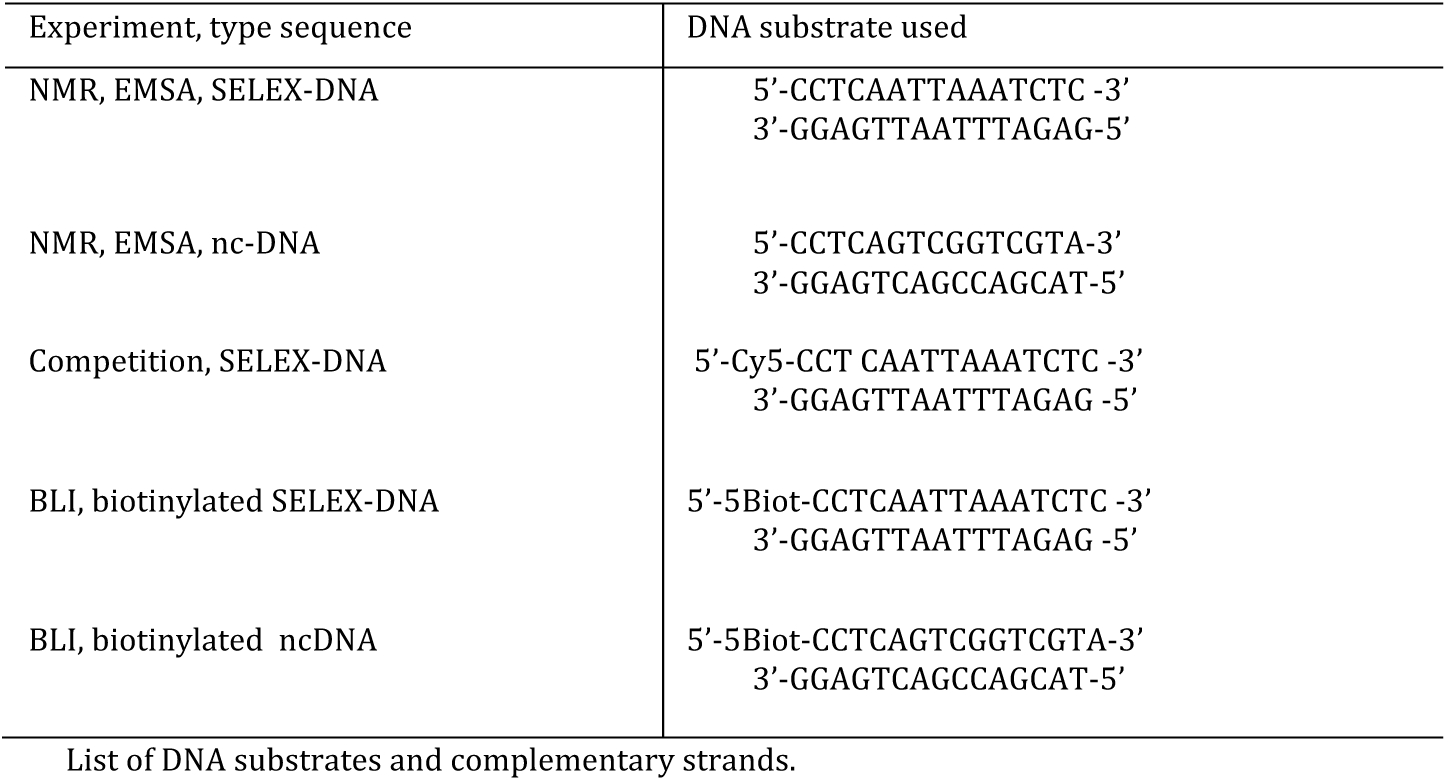

### NMR Data Collection and Analysis

Spectra were collected on a 500 MHz Bruker Avance II, Bruker Avance NEO 600 MHz, or an 800 MHz Bruker Avance II equipped with a cryoprobe. Data processing was performed in NMRPipe and assignments completed in CcpNnmr analysis. Buffer A contains 50 mM potassium phosphate, pH 7.0, 50 mM KCl, 2 mM DTT, 2 mM EDTA in 8% D_2_O.

### NMR Assignments

In order to assign the backbone resonances of BRG1 AT-BD, HNCACB and CBCA(CO)NH, spectra were collected at 500 MHz at 25°C on a sample containing 570 µM of ^15^N-^13^C ATBD in buffer A. Data were processed in NMRPipe and analyzed in CcpNmr analysis. The HNCACB was collected using 40 scans and 80 and 86 complex points in the ^15^N and ^13^C dimensions respectively. For the CBCA(CO)NH, data were collected using 32 scans and 80 and 80 complex points in the ^15^N and ^13^C dimensions. For side chain assignments of the AT-BD, H(CCO)NH and CC(CO)NH spectra were collected at 500 MHz at 25 °C using 570 µM ^15^N-^13^C ATBD in buffer A, and an HCCH-TOCSY was collected at 600 MHz at 25°C. The H(CCO)NH was collected using 32 scans and 70 and 110 complex points in the ^15^N and ^1^H dimensions respectively. The CC(CO)NH was collected using 48 scans and 70 and 68 complex points in the ^15^N and ^13^C dimensions respectively. The HCCH-TOCSY was collected using 32 scans and 80 and 84 complex points in the ^1^H and ^13^C dimensions respectively. The HBHACONH was collected using a sample containing 900 µM ^15^N-^13^C ATBD in the same buffer at 600 MHz at 25°C, using 48 scans and 60 and 100 complex points in the ^15^N and ^1^H dimensions respectively. To determine the side-chain assignments in the DNA-bound state, chemical shifts were tracked upon titration of SELEX-DNA in a ^13^C-HMQC. Sequential ^13^C-HMQC spectra were collected at 800 MHz at 25°C on a 200 µM sample of ^15^N-^13^C ATBD in buffer prepared in 99.8% D_2_O, 50 mM potassium phosphate pH 7.0, 50 mM KCl, and increasing amounts of SELEX-DNA at protein-DNA ratios of 1:0, 1:0.1, 1:0.3, 1:0.5, 1:0.9, 1:1.65. Data were processed in NMRPipe, analyzed in CcpNmr analysis.

To assign the non-exchangeable protons of the SELEX-DNA, homonuclear TOCSY and homonuclear NOESY spectra were collected at 800 MHz at 25°C using 3.2 mM double stranded SELEX-DNA in buffer containing 99.8% D_2_O, 50 mM potassium phosphate, 50 mM KCl, 2 mM EDTA. The propagation of non-exchangeable proton assignments from the apo to the fully saturated DNA was carried out by collecting four spectra using w1 ^12^C-filtered, w2 ^12^C-filtered experiments at 800 MHz at 25 °C using 4 different samples in the same buffer as described above. Each sample contained 1.2 mM of unlabeled SELEX-DNA as apo or in complex with ^15^N-^13^C ATBD. The ratios of DNA:Protein were 1:0, 1:0.15, 1:0.5, 1:1.65.

To assign the exchangeable protons of DNA, along with the H2 of adenines, two homonuclear NOESY spectra were collected encompassing the imino region of DNA at 800 MHz at 25 °C using 5 mM of SELEX-DNA on aqueous buffer (H_2_O) with 50 mM potassium buffer, 50 mM KCl, 2 mM EDTA.

### NMR titrations

Titrations of AT-BD or BD with DNA were carried out by collecting sequential ^1^H,^15^N-HSQC spectra on ^15^N-AT-BD or ^15^N-BD upon addition of increasing concentrations of SELEX-DNA or nc-DNA. Protein and DNA samples were prepared in buffer A. Spectra were collected at 800 MHz at 25°C on 100µM AT-BD at protein:DNA ratios of 1:0, 1:0.1, 1:0.3, 1:0.5, 1:0.9, 1:1.5, 1:2.1, 1:2.7 for SELEX-DNA and 1:0, 1:0.1, 1:0.3, 1:0.5, 1:1.2, 1:1.9, 1:2.8, 1:3.7 for nc-DNA. For the titration into BD alone, spectra were collected at 800 MHz at 25°C on a 100 µM sample of ^15^N-BD at protein:DNA ratios of 1:0, 1:0.3, 1:0.9, 1:1.5, 1:3.3, 1:6.6, 1:13.2 for SELEX-DNA and 1:0, 1:0.3, 1:0.9, 1:1.5, 1:3.3 1:6.6, 1:13.2 for nc-DNA. Data were processed in NMRPipe and analyzed in CcpNmr analysis. The normalized chemical shift perturbations (CSPs or Δδ) were calculated as:

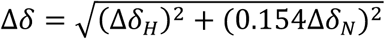

where Δδ_H_ and Δδ_N_ are the change in the ^1^H and ^15^N chemical shift respectively between the apo state and the titration point.

To monitor the binding of DNA to ATBD, a titration was performed by collecting 1D proton spectra of the imino region of unlabeled SELEX-DNA were collected at 800 MHz at 25°C in the presence of increasing amounts of unlabeled AT-BD. Protein and DNA were prepared in same buffer containing 50 mM potassium phosphate pH 7.0, 50 mM KCl, 2 mM EDTA, 2mM DTT. The ratios of DNA:protein were 1:0, 1:0.5, 1:0.1, 1:0.2, 1:0.3, 1:0.5, 1:0.9, 1:1.3, 1:1.7, 1:2.1, 1:2.5, 1:3.3.

### Intermolecular NOESY

To identify the binding interface between the AT-BD and SELEX-DNA an intermolecular NOESY was collected at 800 MHz at 25°C on a 580 µM sample of ^15^N-^13^C ATBD in the presence of 2.2 mM SELEX-DNA. The NOESY was collected in 99.8% D_2_O and 50 mM potassium phosphate, 50 mM KCl, 2 mM EDTA and 2 mM DTT. NOEs between the ^13^C,^15^N-labeled AT-BD and unlabeled SELEX-DNA were detected using ^13^C-edited and ^12^C-filtered 3D NOESY experiments with a mixing time of 150 milliseconds [25].

### Biolayer Interferometry (BLI)

BLI experiments were carried out by tethering double stranded SELEX-DNA or nc-DNA biotinylated at the 5’ end of the forward strand (15 bp each) on a streptavidin sensor. Interference was assayed against increasing concentrations of either AT-BD as analyte in solution. Experiments were carried out in triplicate using an Octet RED96 Biolayer Interferometry (BLI) System (Pall ForteBio, Menlo Park, CA) at 26°C shaking at 1000 RPM. Black 96 well microplates (Greiner bio-one Germany) and Streptavidin sensors (SA, Dip and Read biosensors, ForteBio, California) were used for all experiments. Protein and DNA samples were prepared in 50 mM potassium phosphate, pH 7.0, 50 mM KCl, 0.04% Tween20 and 2 mM EDTA. Each well contained a 200 µL sample, and all probes were pre-soaked in buffer for 45 minutes before each experiment.

The binding assay consisted of a temperature pre-equilibration for 10 min at 26°C, followed by buffer equilibration step of all probes in buffer for 300 s, loading of the biotinylated DNA (50 nM) for 300 s, followed by a baseline step in buffer for 300 s. Following baseline, association was assessed against either 0.25µM, 0.5µM, 1µM, 2µM, 4µM, 8µM, or 15µM ATBD or BD for 300s followed by a dissociation step in buffer for 600 s. Each experiment had a reference well (to correct for buffer drift). Initial trials were performed using a reference sensor to subtract non-specific binding, but no significant non-specific binding was detected. Data was collected using a data acquisition rate of 10 Hz. Steady state analysis was used to determine K_d_ values. Data fitting was performed using the ForteBio Data Analysis 9.0 software. During processing the data, reference wells were subtracted, and data were aligned to the last 10 s of the baseline step, inter-step correction was aligned to dissociation, and Savitzky-Golay filtering was also applied. The fitting was performed by choosing global fitting to R equilibrium option in a 1:1 binding mode which plots the response at equilibrium as a function of the analyte concentration using a single binding site model.

### EMSA and Competition Assay

Electrophoretic Mobility Assays (EMSAs) were carried out on SELEX-DNA or nc-DNA in the presence of increasing amounts of AT-BD. Samples were run on a 10% 75:1 acrylamide:bisacrylamide gel. Gels were casted and run in 0.2X TBE buffer, for 50 minutes at constant 125 V at 4°C. The sample load in each well was 5 µL from a binding reaction containing a volume of 10 µL with 20 mM potassium phosphate, pH 7.0, 80 mM KCl, 2mM EDTA, 2mM DTT. The final concentration of DNA in each well was 53 nM and increasing protein concentrations of 0.04875µM, 0.0975µM, 0.195µM, 0.39µM, 0.781µM, 1.56µM, 3.125µM, 6.25µM, 12.5µM, 25µM, 50µM, and 100µM. Staining of the gel was performed with SYBR Green (ThermoFisher Scientific, Waltham, MA). Visualization of gels was performed in an ImageQuant LAS 4000 imager.

Competition Assays were performed on a 10% 19:1 acrylamide:bisacrylamide gel. Gels were cast and run in 0.2X TBE buffer, for 50 minutes at constant 125 V at 4 °C. Each lane was loaded with 5 µL of a 10 µL binding reaction containing a preformed complex with 0.781µM of BRG1 ATBD in 25 mM potassium phosphate, pH 7.0, 25 mM KCl, 2mM EDTA, 2mM DTT, and 10nM of cy5-labeled SELEX-DNA and using unlabeled SELEX-DNA or nc-DNA as competitor in increasing concentrations of 0.05µM, 0.2µM, 0.4µM, 0.6µM, 1.0µM, 2.0µM, 4.0µM, 8.0µM, 12.0µM, 15.0 µM, 40.0µM. Visualization of gels was performed in an ImageQuant LAS 4000 imager with the Cy5 fluorescence option.

### HADDOCK

The crystallographic structure of the BRG1 BD was obtained from the Protein Data Bank, entry 2GRC [13]. Missing residues and the AT-hook motif KKQKKRGRPPAEKLS were reconstructed *in silico* by using the Modeller program [26] in the UCSF Chimera suite [27]. The DNA sequence 5’-CCTCAATTAAATCTC – 3’ was built using 3D-DART [28].

Molecular docking of the BRG1 AT-BD to SELEX-DNA was carried out with the HADDOCK webserver [29]. The active residues chosen to bias the docking were located in the ZA loop, AB loop, N-terminal, and the complete AT-hook, which were chosen based chemical shift perturbations. All DNA base pairs were considered as active residues to allow the AT-BD system to sample all possible likely conformations along the linear DNA sequence. The docking process consisted of 100 cycles of rigid body energy minimization (EM), semi-flexible simulated annealing, and explicit solvent refinement. In particular, during the rigid-body EM 1000 structures were generated and subsequently reduced to 200 during the semi-flexible and water refinement steps. The top ten docking conformations as determined by their HADDOCK score were used for MD simulations.

### Molecular Dynamics Simulations

For each of the 10 AT-BD/SELEX-DNA complex conformations selected from the HADDOCK docking, three MD simulations were performed. The simulation package GROMACS 2016.4 [30] was used in combination with the AMBER14SB and BSC1[31]force fields. In each simulation, the complex was solvated in a cubic box with a minimum distance of 10 Az to the nearest box edge and surrounded by approximately 15000 water molecules described by the TIP3P water model [32]. System charges were neutralized and Na^+^ and Cl^-^ ions were added to reach 150 mM salt concentration. Systems were minimized for 1000 steps with the steepest descendent method. Complexes were then equilibrated for 100 ps at constant volume and temperature and for 1 ns at constant pressure and temperature. Production simulations were carried out for 100 ns in the NPT ensemble, using a Parrinello-Rahman barostat [33] with a time constant of 2.0 ps to control the pressure and a V-rescale thermostat with a time constant 0.1 ps. Electrostatic interactions were treated with the Particle-Mesh Ewald (PME) method [34, 35] and 9 Az cut-off. An integration time step of 2 fs was used.

Among the conformations obtained through MD simulations, five were selected for further analysis based on how closely they reproduced experimental NOE distance restraints. The systems were refined based on these distances by the addition of distance restraints between the DNA and the AT-BD. This was done in three stages, with the first being a 10 ns simulation in the NVT ensemble and a distance restraint of 1000 kJ mol^-1^ nm^-2^ between Arg1443 NH2 and 1A5 N1, the second a 10 ns NPT simulation with distance restraints of 1000 kJ mol^-1^ nm^-2^ between Arg1443 CD and 1A9 C2, Arg1443 CD and 2A8 C2, Arg1445 CD and 1A6, Arg1445 CD and 2A9 C2, and the third a 10 ns NPT simulation with additional restraints between Leu1488 CD1 and 2T7 C7, Tyr1498 CE1 and 2T7 C7. Following these three restrained simulation stages, 500 ns of unrestrained simulations were performed.

Structure equilibrations were evaluated by analyzing the root mean square deviations (RMSD). In the calculation of atomic RMSD protein rotation and translation were eliminated by a mass-weighted least squares fit with respect to the reference starting structure. VMD [36] was used for analysis of the hydrogen bonds and for trajectory visualization. In particular, hydrogen bonds were calculated between residue pairs with occupancies computed as the percent of MD frames with bond donor-acceptor distance and angle cutoffs of less than 3.5 Az and 30 degrees. Contact frequency maps were calculated for every frame of the equilibrated portion of the trajectories with GROMACS mdmat (trunc distance 4.2 Az). Do_x3DNA and dnaMD tools [37, 38] were used to analyze global and local DNA properties during MD simulation.

## Results

### BRG1 AT-BD has preference for a 10-basepair AT-rich sequence

We previously found that the BRG1 BD can associate with DNA, and that the BD and adjacent AT-hook act as a composite domain to bind DNA in a multivalent manner [15]. Though we found a preference for AT-containing DNA, only two DNA sequences were tested in that seminal work. We therefore performed <u>S</u>ystematic <u>E</u>volution of <u>L</u>igands by <u>Ex</u>ponential enrichment followed by deep <u>seq</u>uencing (SELEX-seq, Figure 1B) [39] to measure the sequence preference of AT-BD in a comprehensive and unbiased fashion.

Having previously observed that the AT-BD occludes a single helical turn of double stranded (ds) DNA (∼10bp), we sought to determine whether this composite domain has a preference for sequences of this length or longer [15]. Accordingly, we incubated AT-BD with a randomized 15 base pair library and separated bound from free by electrophoretic mobility shift assay (EMSA). The bound sequences were isolated by excising the shifted band from the gel, amplifying the sequences, and repeating twice, for a total of three rounds of enrichment (Figure 1B). Using the SELEX package [39], we found that information gain reached a maximum with a binding site of 10bp (Supplementary Figure S1A), providing strong evidence that the 10bp footprint is accurate. We then used the SelexGLM package [24] to calculate a position-specific affinity matrix over a 10bp footprint, which was converted to logo format using MATRIXReduce [40]. The most favorable motif is composed of an asymmetric AT-rich core 8bp sequence (AATTAAAT) (Figure 1B, right, Supplementary Figure S1C), with slight preference for G at the first and C at the last position. Interestingly, the AT-BD appears able to accommodate either As or Ts, with little penalty for substitution of one over the other within the core motif, except at position 7.

To validate the preference for the SELEX-determined sequence we carried out electromobility shift assays (EMSAs) with the AT-BD and 15 bp double-stranded DNA containing either a non-consensus sequence (nc-DNA: 5’-CCTC<u>AGTCGGTCG</u>TA-3’) or the SELEX determined consensus sequence (SELEX-DNA: 5’-CCTC<u>AATTAAATC</u>TC - 3’). Binding of AT-BD to SELEX-DNA led to discretely shifted bands for the complex at lower AT-BD concentrations (Supplementary Figure S1B, see lanes 5 and 6) with less discrete higher-order complexes evident at higher AT-BD concentrations. In contrast, EMSAs for nc-DNA:AT-BD complexes ran as smears for all concentrations tested, reflecting less stable complexes (Supplementary Figure S1B). In addition, we performed competition assays using Cy5-labeled SELEX-DNA (Figure 1C), and either unlabeled SELEX-DNA or unlabeled nc-DNA as a competitor. Titration of the unlabeled SELEX-DNA competitor led to complete disappearance of the Cy5-SELEX-DNA-AT-BD complex (Figure 1C, top) whereas titration of nc-DNA was not able to efficiently compete off the Cy5-labeled SELEX-DNA (Figure 1C, bottom). Together this indicates a higher affinity, more stable complex with SELEX-DNA relative to nc-DNA.

To quantify the increase in affinity for the preferred sequence we utilized biolayer interferometry (BLI) (Figure 1D). Fast kinetics of binding precluded reliable fitting of kinetic parameters, however consistent K_d_s were obtained through steady state analysis of the level of response as a function of AT-BD concentration. This yielded K_d_=4.9μM for nc-DNA and K_d_=1.8μM for SELEX-DNA, or a ∼3-fold greater affinity for the SELEX-DNA than for the non-consensus DNA sequence (Figure 1D).

### Chemical shift mapping reveals a unique binding mode to SELEX-DNA

To gain insight in the molecular basis of complex formation with SELEX-DNA we utilized NMR spectroscopy [41–44]. The complex was initially examined through chemical shift perturbation (CSP) experiments. Sequential ^1^H,^15^N-HSQC spectra were collected on ^15^N-AT-BD while titrating in either nc-DNA or SELEX-DNA. Both substrates led to substantial CSPs indicating binding (Supplementary Figure S2). Both titrations were in fast-to-intermediate exchange on the NMR timescale consistent with micromolar affinity. Notably, peaks corresponding to the BD and the AT-hook shifted concomitantly (i.e. in response to the same concentration of DNA), suggesting a multivalent contribution from both the BD and AT-hook to DNA binding, consistent with our previous observations [15, 45].

Backbone resonances were assigned using standard triple resonance experiments (Supplementary Figure S3A)[46, 47]. Plotting the normalized CSPs against residue number for both titrations reveals the residues that are directly and indirectly involved in binding to nc-DNA or SELEX-DNA (Figure 2A). In both titrations, we observed significant perturbations in residues corresponding to both the AT-hook and the BD. However, SELEX-DNA also induced significant CSPs in linker residues Lys1450, Leu1451, Ser1452, and Asn1454 that were not seen with nc-DNA. SELEX-DNA also led to substantially larger perturbations compared to nc-DNA for residues in the AT-hook (Lys1438, Lys1439, Gln1440 and Lys1442) and residues in the BD (Ser1490, Lys1492, Ile1501, Arg1502, Lys1509, His1517, Tyr1519 and Arg1520) (Figure 2A, bottom). Mapping the CSPs onto the previously solved structure, reveals a binding pocket which is formed by *α*A and the ZA loop for both nc-DNA and SELEX-DNA, with additional contributions by the very N-terminal end of *α*Z and the AB loops for SELEX-DNA (Figure 2B). This binding pocket is lined with basic residues that form a larger basic patch on the BD surface (Supplemental Figure S3B), consistent with what we previously observed [15]. Together, this data suggests a more extensive binding pocket and a more stable complex with SELEX-DNA compared to nc-DNA.

**Figure 2.**
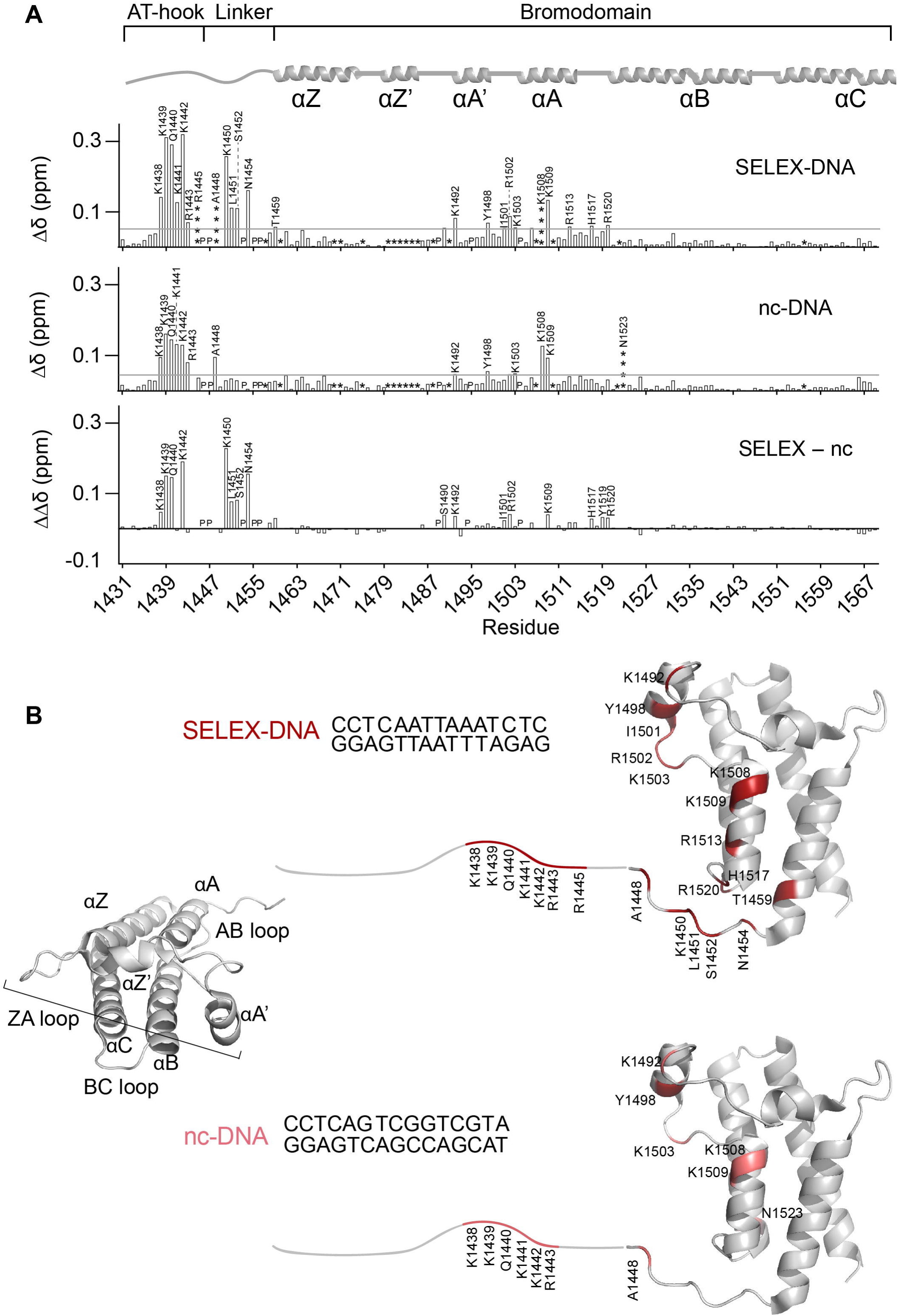
NMR reveals the molecular basis of enhanced binding to SELEX-DNA. (A) Plots of normalized CSPs (Δδ) for AT-BD with SELEX-DNA (top), nc-DNA (middle), and the difference between the two (ΔΔδ, bottom). Residues perturbed more than the average plus two standard deviations (denoted by the gray line) are labeled. Where the average was calculated after trimming the top 10% of perturbations. “P” indicates a proline, “*” indicates an unassigned residue and “****” represent a signal that broadened beyond detection upon titration with ligand. The AT-BD secondary structure architecture is depicted on top of the plots for reference. (B) Residues in the AT-BD with significant CSPs upon binding to the SELEX-DNA (top) or nc-DNA (bottom) are plotted onto a cartoon representation of the previously solved structure of the BD (PDB ID 3UVD), with the AT-hook drawn in (a small break is left between the structure and the drawn-in segment).

In addition to changes in the magnitude of CSPs between the two substrates, we investigated differences in the trajectory of CSPs induced by either SELEX-DNA or nc-DNA. Differences in the trajectory of CSPs were observed in several residues in the AT-hook, linker, and the BD indicating a unique binding mode. These included residues Lys1439, Lys1441, Arg1443, Lys1450 and Leu1451 in the AT-hook and linker and residues Thr1459, Tyr1498, and Tyr1519 in the BD (Figure 3).

**Figure 3.**
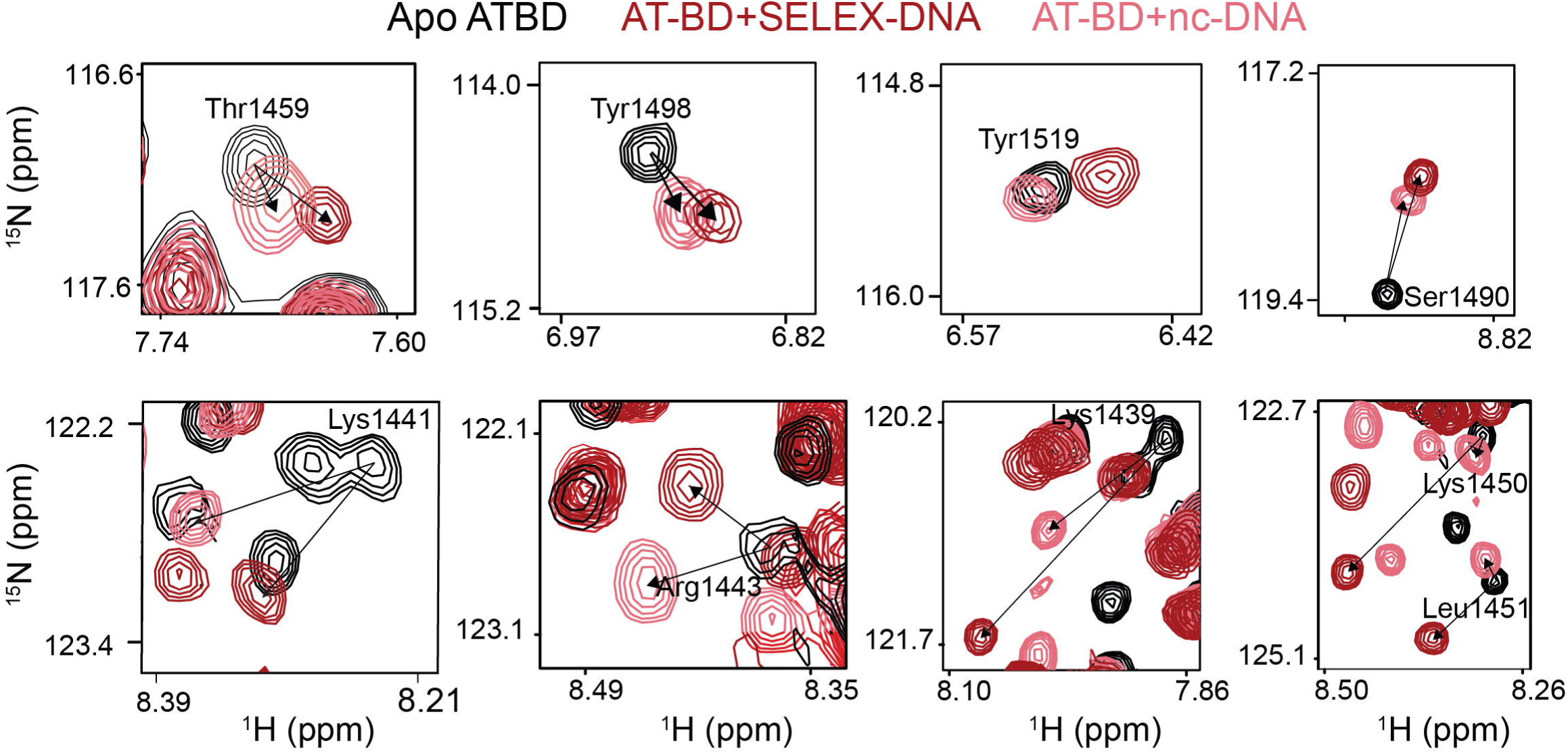
NMR reveals unique binding to SELEX-DNA. ^1^H,^15^N-HSQC spectral overlays are shown for AT-BD residues saturated with either SELEX-DNA (red) or nc-DNA (salmon). Shown are resonances for residues which show a unique trajectory of CSP between the two substrates, indicating a unique binding mode. This was observed for residues in the bromodomain (Thr1459, Tyr1498, Tyr1519, Ser1490) the AT-hook (Lys1439, Lys1441, Lys1443) and the linker (Lys1450, Lys1451).

A similar analysis was carried out for the BD alone. Spectra of the ^15^N-BD construct saturated with nc-DNA or SELEX-DNA were overlaid. As was seen with the AT-BD, larger CSPs were induced in the BD resonances by SELEX-DNA (Supplementary Figure S4). However, in contrast to the AT-BD there were no perceptible changes in trajectories of CSPs between nc-DNA or SELEX-DNA (Supplementary Figure S4). Together the CSP experiments suggest that both the AT-hook and BD contribute to the preference for SELEX-DNA. The increased magnitude of CSPs indicates that both the BD alone and AT-BD form a more stable complex with SELEX-DNA compared to nc-DNA. In addition, the additional CSPs and altered trajectories indicate that the AT-BD adopts a unique structure in binding to SELEX-DNA as compared to nc-DNA.

### The AT-hook and BD span the minor and major grooves of SELEX-DNA

Having defined the DNA-binding interface of the protein, we sought to identify the protein-binding interface on DNA, also using NMR spectroscopy. Resonances for SELEX-DNA non-exchangeable (H2, H5, H6/H8, H1’, H2’, H2’’) and exchangeable (iminos) protons were assigned (Supplementary Figure S5). Initial collection of 1D-proton spectra of SELEX-DNA upon titration of increasing amounts of AT-BD revealed CSPs indicating binding (Figure 4A, Supplementary Figure S6A). Notably, the most substantial CSPs were seen for the imino protons of A6, T7, T8, A9 and A10, which are in the core of the SELEX determined sequence.

**Figure 4.**
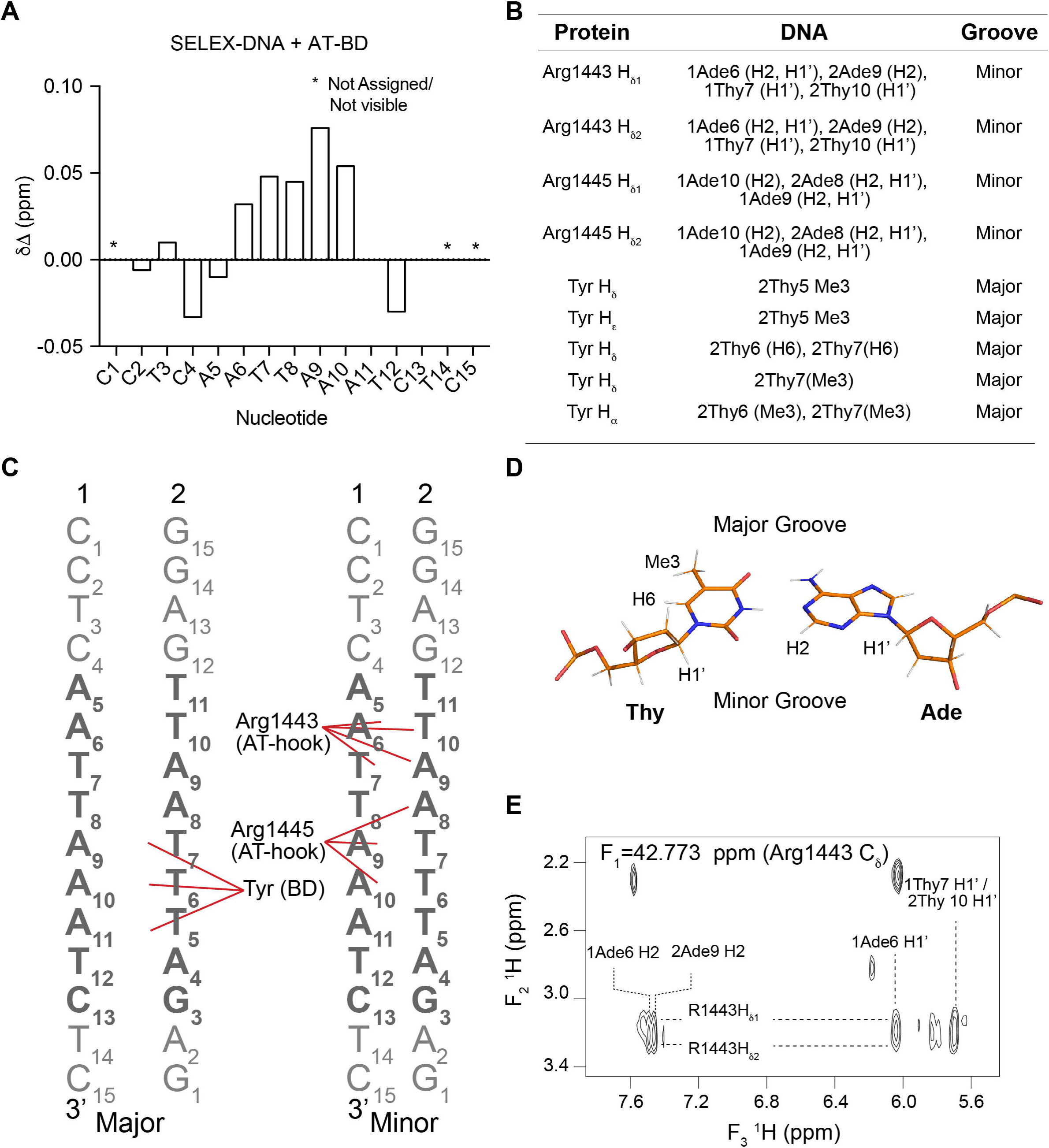
The AT-BD spans the major and minor groove. (A) CSPs (Δδ) of the imino resonances of the SELEX-DNA saturated with AT-BD. “*” indicates an unassigned or not visible resonance. (B) Table of Intermolecular NOEs observed between the ^13^C-AT-BD and ^12^C-SELEX-DNA. (C) Schematic of AT-hook and BD contacts with SELEX-DNA (strands 1 and 2 labeled) in the major (left) and minor (right) grooves. Contacts are highlighted by red lines. The core SELEX determined sequence is in bold. (D) structure of an AT base pair with hydrogens forming NOEs labeled, as well as the major and minor groove sides. (D) Representative NOE cross-peaks between AT-BD and SELEX-DNA indicate that these protons are <6Å from each other in space.

We then measured intermolecular NOEs to detect proton-proton distances of <6 Å between the AT-BD and SELEX-DNA via an isotope edited/filtered approach using ^13^C/^15^N-labeled AT-BD and unlabeled SELEX-DNA. In this experiment, the intermolecular NOE cross-peaks between ^13^C-H protons of AT-BD and ^12^C-H protons of DNA were detected. Side chain assignments of the AT-BD were made by tracking the chemical shift in a ^1^H,^13^C-HMQC upon titration of SELEX-DNA (Supplementary Figure S6B).

Several intermolecular NOEs were observed between the AT-hook and adenines in SELEX-DNA sequence (Figure 4 and Supplementary Figure S6C). Specifically, NOEs were seen between the H*δ* proton of Arg1443 in the AT-hook and the H2 protons of both 1A6 and 2A9 in SELEX-DNA, where 1 and 2 refer to the forward and reverse strand respectively (Figure 4B,C,E). Similarly, we observed NOEs between Arg1445 H*δ* proton of the AT-hook and the H2 protons of 1A9 and 2A8. Consistent with this, we also observed NOEs between Arg1443 H*δ* protons and H1*′* protons of both 1T7 and 2T10 and between Arg1445 H*δ* protons the H1*′* protons of 1A9 and 2A8 (Figure 4B,C and Supplementary Figure S6C). In B-form DNA, H1*′* protons and adenine H2 protons are most easily accessible through the minor groove, indicating that the AT-hook is inserting into the minor groove (Figure 4D) [19, 48]. Notably, in ^1^H,^13^C-HMQC spectra of AT-BD single peaks are observed for the H*δ* protons of Arg1443 and Arg1445, but upon titration of SELEX-DNA these peaks split into two. This suggests a fixed position of these arginines upon binding to DNA (Supplementary Figure S6B). This is consistent with a canonical AT-hook binding mode, in which the narrowed minor groove of AT-rich DNA is favored. Indeed, evidence of minor groove narrowing in the apo SELEX-DNA is supported by an inter-strand NOE observed in the homonuclear NOESY spectrum between 2T7 H1’ and 1A10 H2 protons (Supplementary Figure S5F) [49, 50].

In addition, NOEs were observed between the BD and SELEX-DNA (Figure 4 and Supplementary Figure S6C). These NOE cross-peaks were observed to the methyl groups of 2T5, 2T6, and 2T7, indicating that the BD is interacting in the major groove (Figure 4B,C,E and Supplementary Figure S6C). The corresponding protein chemical shifts for these NOE cross-peaks are consistent with H*δ*/H*ε* protons of two tyrosine residues. Unfortunately, these tyrosines were not observed in either the HCCONH nor the CCONH experiments, and thus they could not be definitely assigned. However, only two tyrosines experienced significant CSPs in the ^15^N-HSQC upon titration of SELEX-DNA into AT-BD, and only three tyrosines are in the binding pocket. These are Tyr1497 and Tyr1498 in the ZA loop and Tyr1519 at the end of the αA helix (see Figure 2). This strongly suggests that two of these three tyrosine residues are making contact with the SELEX-DNA major groove, and thus at least one must be a Tyr in the ZA-loop. Though these tyrosines could not be assigned, given the positioning of the AT-hook, one tyrosine should be adjacent to 2T6 and 2T7, and another should be adjacent to 2T5. An additional NOE was seen between a tyrosine H*δ* proton and the H6 protons of 2T6 and 2T7 (Figure 4B,C and Supplementary Figure S6C).

Together, the NOEs reveal that the AT-hook and BD span the minor and major grooves [51]. The NOEs are consistent with the Arg-Gly-Arg (Arg1443, Gly1444, Arg1445) of the AT-hook inserting into the minor groove of the narrowed AT-rich segment. In addition, the NOEs between the BD and DNA would be consistent with two aromatic/methyl interactions between tyrosines and thymines, a common major groove mode of binding. At least one of these interactions is being formed with the ZA loop with an additional interaction in the ZA loop or αA helix. The NOEs are consistent with the binding pocket identified by CSPs (Figure 2). The CSPs suggest further electrostatic interactions between basic residues in both the AT-hook and BD and SELEX-DNA. This is supported by the detection of several additional NOEs that would be consistent with Lys/DNA interactions, however substantial spectral overlap precluded our ability to resolve these.

### Molecular modeling reveals the importance of the linker in AT-BD binding

To better understand how the AT-BD docks and binds SELEX-DNA, a hierarchy of computational methods were used to model the complex. Initial complexes of AT-BD with SELEX-DNA were obtained using the data-driven docking program HADDOCK, which is a flexible docking approach that imposes distance restraints based on experimentally defined binding interfaces. Unambiguous interaction restraints on the AT-BD were assigned based on CSP analysis (see Figure 2), while no restraints were imposed on the DNA. The structures with the top-ten highest HADDOCK Z-score were selected for further analysis. These poses were used as starting structures for triplicate all-atom molecular dynamics (MD) simulations of 100 ns in length.

Several binding modes of the AT-BD along the SELEX-DNA were obtained in the initial ten docked structures, and the observed binding modes observed at the end of the 100 ns simulations were similar to their starting conformations. Of these, five had the AT-hook bound to the SELEX-DNA minor groove and the BD in proximity of the major groove consistent with the NOE data. In the other conformations the AT-hook was bound to the DNA major groove with αA in either the major or minor groove, or the AT-hook bound in the minor groove, but inconsistent with the experimental data. Notably, these were also predicted to be the least favorable based on docking score. As such, these five binding modes were discarded from further analysis, reducing the number of binding conformations from ten to five.

MD simulations were rerun on the remaining five conformations using a multi-step restraint-enabled refinement approach, followed by 500 ns of unrestrained MD simulations (Supplementary Figure S7A). In this step the restraints were based on experimental NOEs. At the end of these runs, three of the simulations had the AT-hook consistently bound in the DNA minor groove while in the other two it was associated in the major groove. The two binding modes with the AT-hook bound to the major groove were discarded and further analysis was performed on the three states with the AT-hook bound to the minor groove. Notably, two of the three are not fully consistent with the NMR data. In binding mode 2 the AT-hook is associated with, but not inserted into the minor groove, while in binding mode 3 it is positioned in a reverse manner in the minor groove (Supplementary Figure S7). The position of the AT-hook and BD in binding mode 1 however, is overall consistent with the experimental data (Figure 5A,C).

**Figure 5.**
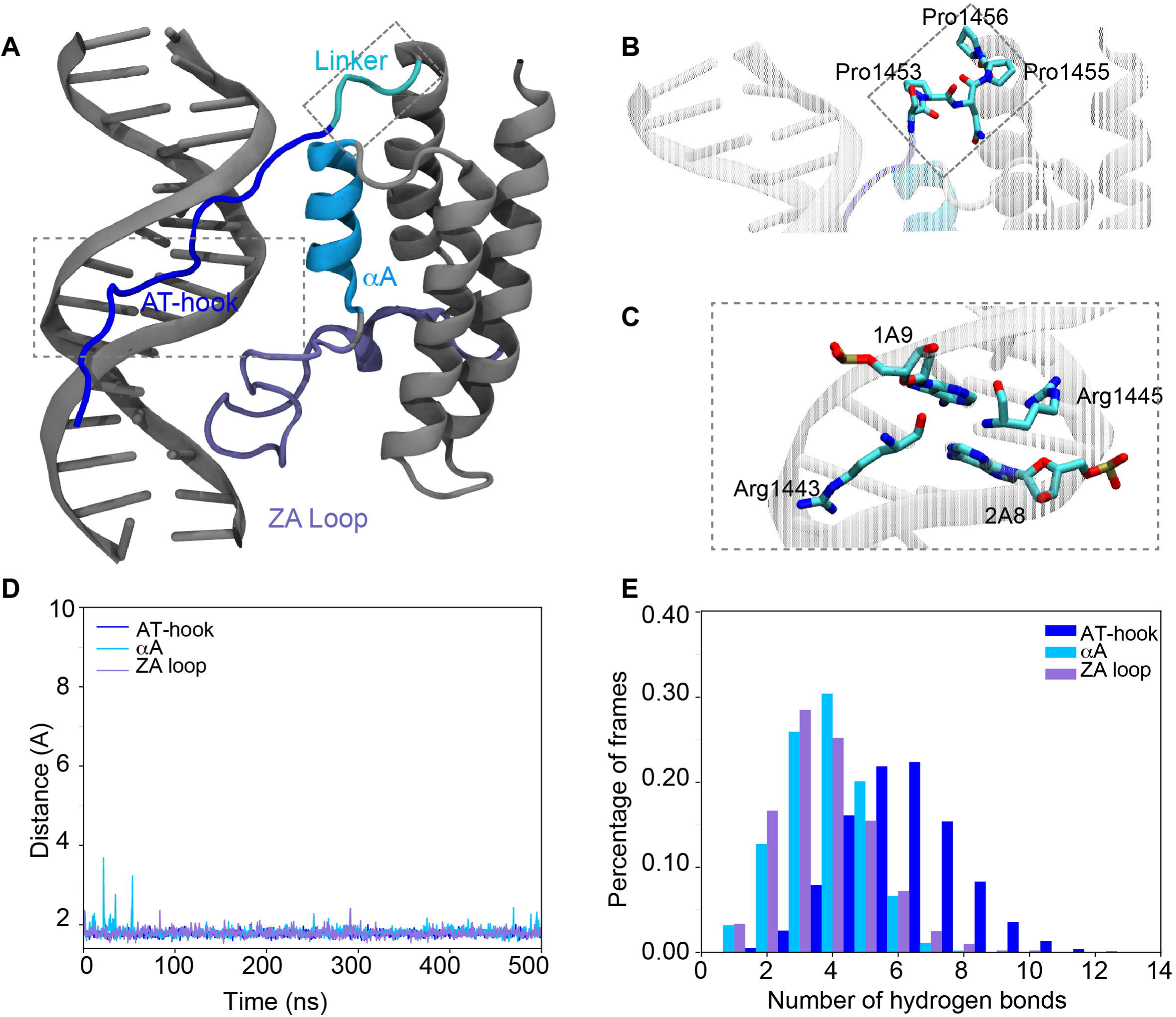
Molecular model of the AT-BD/SELEX-DNA complex. (A) Model of AT-BD/SELEX-DNA complex. The AT-hook is highlighted in blue, the αA in deep sky blue, the ZA loop in ice blue and the linker in cyan. The DNA and the rest of the BRG1 BD are in gray. In this binding mode (mode 1) the AT-hook is positioned in the minor groove and the ZA loop the major groove, consistent with NOE data. (B) A zoom in of the linker shows a significant bend induced at in the polypeptide backbone at Pro1456. (C) A zoom in shows the AT-hook arginines (1443 and 1445) inserted into the minor groove. Carbons are colored in cyan, oxygens in red and nitrogens in blue. (D) The minimum distance between AT-BD elements and SELEX-DNA. (E) The number of hydrogen bonds between AT-BD elements and SELEX-DNA in the percentage of frames during the 500 ns simulations.

The overall stability of the binding modes was assessed by calculating the root mean square deviation (RMSD) of AT-BD relative to SELEX-DNA. The average RMSD for all three binding modes was *≤* 4 Å during the simulations, with the exception of binding mode 2 in which there was an increase in the final 50 ns that corresponded to a slight twisting of αA relative to the DNA (Supplementary Figure S7B). However, only in binding mode 1 are both the AT-hook and BD associated stably with the DNA throughout the simulation.

To further investigate the binding mode, minimum distances between the DNA and AT-BD elements were calculated for each binding mode. Our results show that the minimum distance between the AT-hook and DNA during the 500 ns simulation (as measured between any protein heavy atom and any DNA heavy atom) is stable at 1.76 ± 0.08 Å for all three binding modes. For the BD, in binding mode 1, the distance between αA and the DNA is stable at 1.81± 0.13 Å and between the ZA loop and DNA is stable at 1.79 ± 0.09 Å over the course of the 500 ns simulations (Figure 5D). In binding modes 2 and 3 the BD is far more dynamic with the DNA/αA distance fluctuating from 1.58 Å to 5.03 Å, and the DNA/ZA loop distance fluctuating from 1.46 Å to 9.55 Å (Supplementary Figure S7C,D). DNA base-pair parameters for both DNA free in solution and bound to the AT-BD were calculated to quantify the effects of binding on the DNA. Results showed little differences in the base-pairs parameters in all simulations, suggesting that AT-BD binding does not change the geometry of the double helix (results not shown).

Contact analysis showed that AT-hook and BD made significant contacts with the DNA in all three simulations. The highest contact frequency is observed for Lys1442, Arg1443, Gly1444 and Ala1448 in the AT-hook consistent with NMR CSPs. Analysis of hydrogen bonding between the DNA and residues in the AT-hook, αA and ZA loop demonstrate that the AT-hook remains stably bound to the DNA. In contrast, the BD interactions are more transient (Figure 5E and Supplementary Figure S7C,D). For binding mode 1 the average number of hydrogen bonds formed between the DNA and AT-hook is the highest, 5.60 ± 1.76, whereas αA and the ZA loop formed fewer hydrogen bonds at 2.76 ± 1.26 and 2.72 ± 1.46 respectively. Similar trends are seen for binding modes 2 and 3 (Supplementary Figure S7C,D).

Notably, the model indicates that the composition of the linker between the AT-hook and BD is critical for spanning the major and minor grooves. From the AT-hook, the linker is seen to wrap around one strand of the SELEX-DNA allowing the BD ZA loop to insert into the major groove. The conformation of Pro1456 in the linker adopts a unique conformation (Figure 5B and Supplementary Figure S7E) and provides a critical turn in the polypeptide necessary for both the AT-hook and BD to associate with both the major and minor grooves. This is consistent with the CSP analysis, in which the linker resonances were uniquely shifted for SELEX-DNA.

Together, simulation results show that the AT-hook and the BD can bind to the DNA in different locations. Though only one of these conformations is fully consistent with the experimental data, the others may be representative of minor populated states. Both minimum distance calculations between the DNA and AT-BD, and hydrogen bond analysis show that the complex is driven by interactions of the AT-hook, αA and ZA loop with DNA. The AT-hook interaction is the most stable while the BD interaction is more transient. Finally, the simulations reveal that the linker is critical in mediating the multivalent interaction of the AT-hook and BD with SELEX-DNA.

### Cancer mutations alter the mode of AT-BD DNA binding

It is estimated that 20% of human tumors present a mutation in at least one of the subunits of BAF [52]. This includes mutations in the DNA binding pocket of the BRG1 AT-BD. We investigated six of these mutations to determine the effect on DNA binding: R1445W in the AT-hook, P1456L in the linker, R1502H and P1504L in the ZA loop, and R1515H in the αA helix. In addition, we examined R1520H in the AB loop. To investigate global DNA association, EMSAs were carried out with the 147 bp Widom 601 DNA naked or formed into unmodified NCPs and each of the six mutant AT-BD constructs (Supplementary Figure S8). Notably, the 601 sequence includes four AT-rich regions. Qualitatively, EMSAs revealed the largest abrogation of binding for mutants P1456L (linker) and R1502H (ZA loop), as assessed by the disappearance of free substrate. Mutants R1515H (αA helix) and R1445W (AT-hook) also show abrogation of binding but to a lesser extent. Notably, almost all mutants show some change in the laddering pattern observed upon binding to the DNA or NCPs, with the exception of P1504L. Similar results were seen for both naked and NCP DNA. Together, this indicates that excepting P1504L, all mutants tested have an effect on the mode of AT-BD DNA binding.

We further investigated P1456L (linker) and R1502H (ZA loop) using NMR spectroscopy and BLI. Binding affinities for SELEX-DNA were determined by BLI and yielded only small changes as compared to wild type (WT): K_d_=1.2 µM±0.1 for P1456L, K_d_=2.4±0.2 µM for R1502H, K_d_= 1.8 µM±0.2 WT. However, changes seen in the EMSAs (Supplementary Figure S8) suggest that the disruption in binding mode leads to more substantial defects on longer pieces of DNA. To assess if either mutation affected complex formation, ^1^H,^15^N-HSQC spectra were collected on both mutants. Comparing the mutant spectra to WT revealed that both mutants retain a similar overall fold (Supplementary Figure S9). To investigate differences in DNA binding by the mutants as compared to WT, sequential ^1^H,^15^N-HSQC spectra were collected on ^15^N-labeled mutants upon addition of increasing amounts of SELEX-DNA. Both mutants show differences in CSPs around the mutation as expected. However, for both mutants, differences are also seen in the magnitude of CSPs throughout the AT-hook and BD (Figure 6) indicating a change in the mode of DNA binding. Notably, these were most pronounced for the linker mutation, which is highly consistent with the model that predicts that the linker composition is critical to allow the AT-hook and BD to span the major and minor grooves.

**Figure 6.**
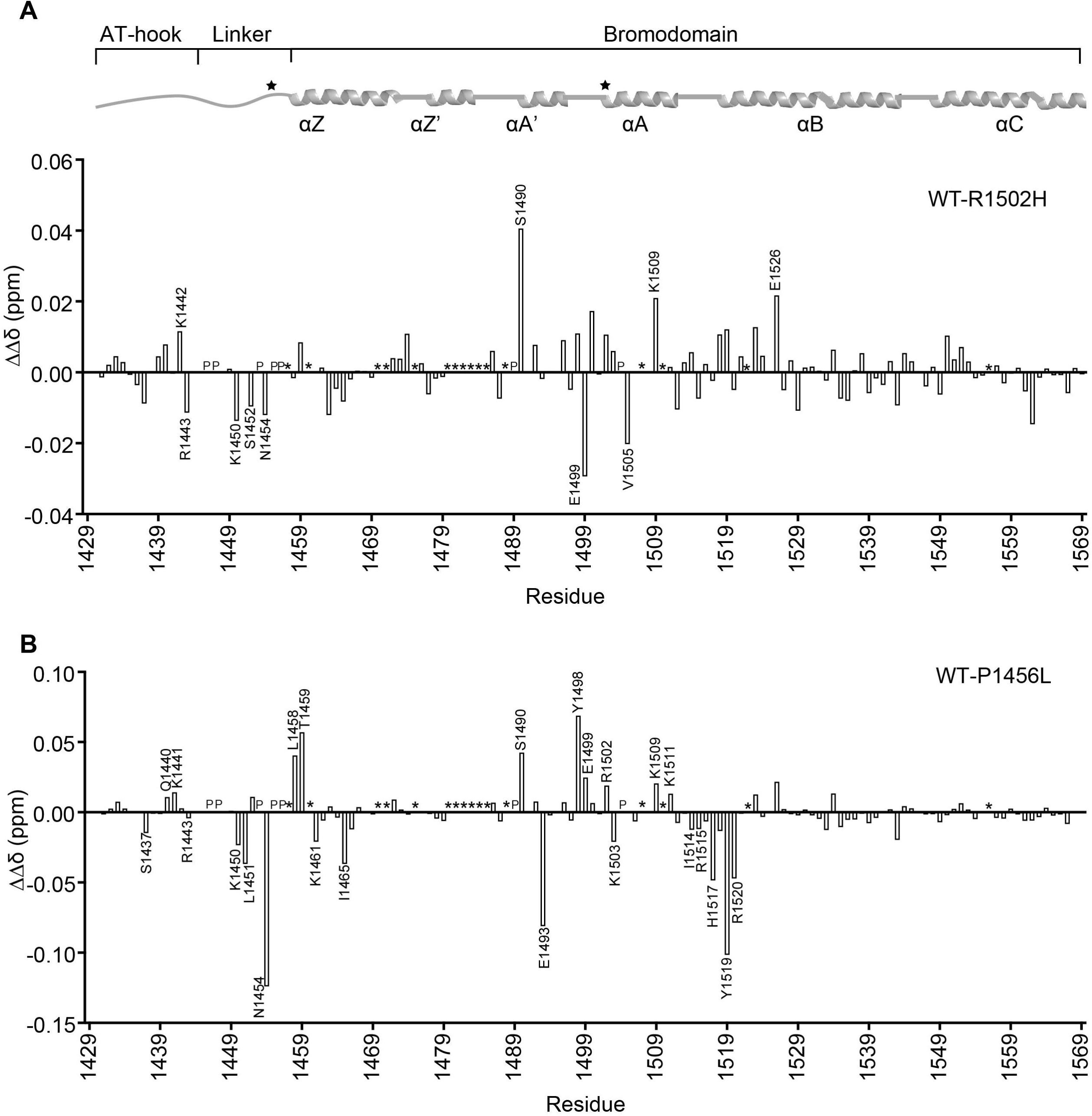
Cancer mutations alter the mode of DNA binding. Differences in the magnitude of CSPs (ΔΔδ) for binding of AT-BD to SELEX-DNA were computed between wild-type (WT) and the R1502H (A) or P1456L (B) mutants. Residues which show substantially different binding are marked. “P” denotes proline and “*” denotes unassigned residues. The AT-BD secondary structure architecture is depicted on top of the plots for reference with mutation sites marked with a star.

## Discussion

Here we have identified a consensus sequence for the AT-hook and bromodomain of BRG1 and have determined the molecular basis for DNA binding and specificity. We show that the AT-hook and bromodomain span the minor and major grooves of the consensus sequence, with the AT-hook inserted into the minor groove and the bromodomain ZA loop inserting into the major groove. This binding mode is reminiscent of HOX proteins, in which a homeodomain inserts into the major groove and the nearby disordered “N-terminal arm”, often arginine rich, associates in the adjacent minor groove [53]. Sequence specificity appears to arise largely from shape recognition of the DNA by the AT-hook with additional base specific contacts by the BD. In particular, the narrowing of the minor groove characteristic of AT-rich sequences leads to electrostatic focusing, which is known to enhance the affinity of arginines for DNA [21, 54]. Thus, the AT-hook has preference for this shape. In addition, two base specific contacts were detected between tyrosines in the BD and thymines in the major groove that help stabilize interaction with the consensus sequence. Notably, the linker composition appears critical for this binding conformation. Indeed mutation of the linker leads to changes in both the AT-hook and BD binding.

AT-hooks are known drivers of the interaction of HMG proteins with DNA [22]. They have also been found in other transcription factors such as ELF3, in which an AT-hook adjacent to the ETS specificity domain is required for function. For BRG1 the AT-hook and bromodomain appear to be structurally and functionally coupled and act in a composite manner to associate specifically with DNA. Here the AT-hook is not simply an accessory motif, neither is it solely defining DNA binding. Notably, there is substantial variation in the target sequence of AT-hooks, likely arising not only from differences in motif sequence, but also from the surrounding protein context.

AT-hooks are found in a number of chromatin associated proteins. Of the 61 human bromodomain-containing proteins, we found that 14 also contain at least one canonical GRP containing AT-hook. Of these, the bromodomain of ASH1L is predicted to bind nucleic acid [15]. In addition, several other reader domains (e.g. chromodomains and PWWP domains) that have been identified to bind nucleic acid contain adjacent canonical AT-hooks, AT-like-hooks, extended AT-hooks, or extensions rich in basic residues [55]. Thus, this multivalent mode of interaction may be common. However, additional studies would need to be done to determine the sequence specificity of these motifs and the molecular basis of binding.

Though we have shown that the AT-BD can associate with DNA formed into the nucleosome particle, we do not yet know the effect of nucleosome structure on the mode of DNA binding. For most transcription factors nucleosome wrapping substantially inhibits association with DNA both by inhibiting association and enhancing dissociation [56, 57]. However, for others, binding to nucleosomes is as efficient as naked DNA, and some even prefer nucleosomal DNA [58, 59]. It is not fully clear how this is achieved mechanistically but it has recently been shown that this is due to decreased dissociation rates [60] and for some may include stabilizing contacts with the histones [61]. Our previous work revealed that concomitant binding of DNA and H3K14ac histone tails by the BD is possible, suggesting that the AT-BD may participate in dual histone/DNA binding at acetylated nucleosomes [15]. Further work will be necessary to determine how this may alter DNA sequence specificity and mode of binding.

It remains to be determined exactly how the AT-BD DNA binding activity contributes to the function of the BAF complex. It may play a role in targeting or retaining BAF at certain regions of chromatin. In addition, it may play a role in the chromatin remodeling activity itself, through regulation of ATPase or remodeling activity. The results presented here provide tremendous insight into the mechanisms of function of this composite domain and lay the groundwork for determining its role in BAF function. In addition, they provide insight into the deleterious effects of cancer mutations.

## Acknowledgements

We would like to thank Lokesh Gakhar for his assistance in experimental design and data analysis, Chris Ptak, Madeline Shea, Adina M. Kilpatrick, and Ryan Mhaling for many thoughtful discussions of data.

