## Supplementary Figures for "The Molecular Basis of Specific DNA Binding by the BRG1 AT-hook and Bromodomain"

**Sanchez et al., Supplementary Figures**

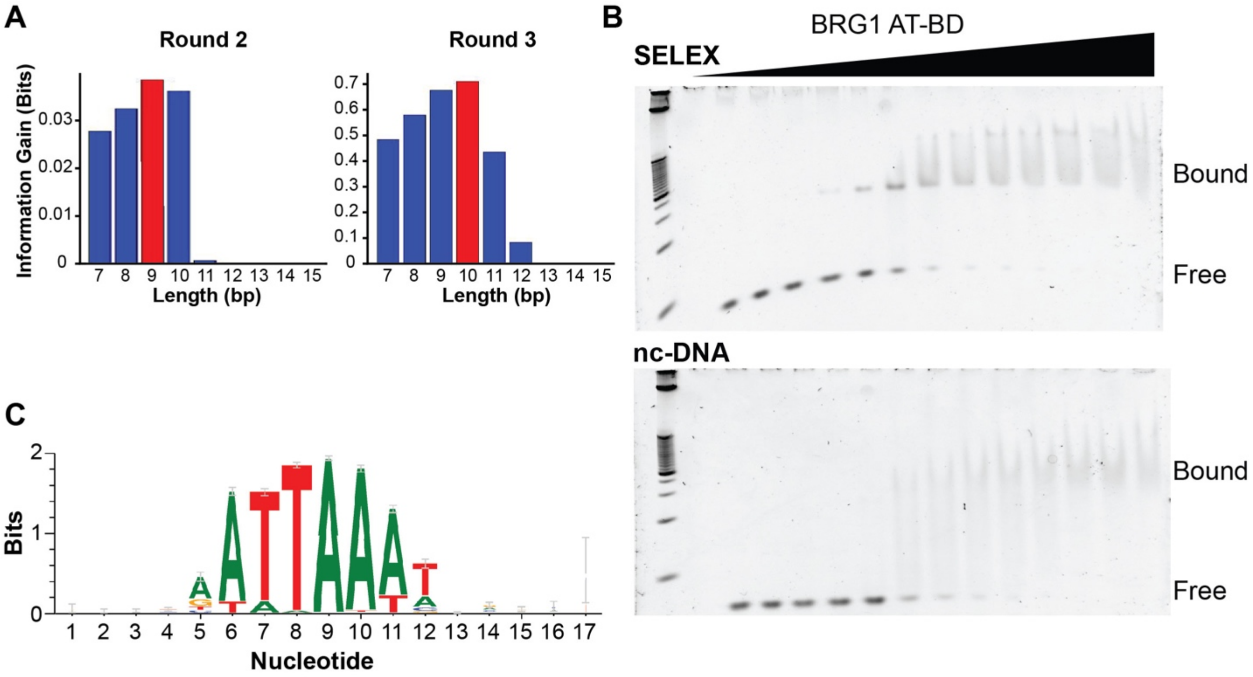


**Figure S1. SELEX-seq defines the length and composition of the AT-BD binding site. (A)** SELEX-seq was performed for a total of 3 rounds. Information gain analysis (SELEX,R/Bioconductor) shows maximal information gain is achieved with a 9bp long site after two rounds (left). After 3 rounds maximal information gain is achieved with a binding site length of 10bp. **(B)** Position weight matrix (PWM) for the 2000 most highly enriched sequences after Round 3 of SELEX-seq. A core 9bp binding site (Nucleotides 4-12) as well as some downstream preference (Nucleotides 14-15) is evident that closely resembles the 10bp Position Specific Affinity Matrix (PSAM) calculated by SelexGLM based on enrichment from Round 2 to Round 3 (Figure 1C). **(C)** Elelctromobility shift assays (EMSAs) assessing the affinity of AT-BD for the consensus motif (SELEX-DNA, top) defined by the SelexGLM PSAM (Figure 1C) and a non-consensus binding site (nc-DNA, bottom).

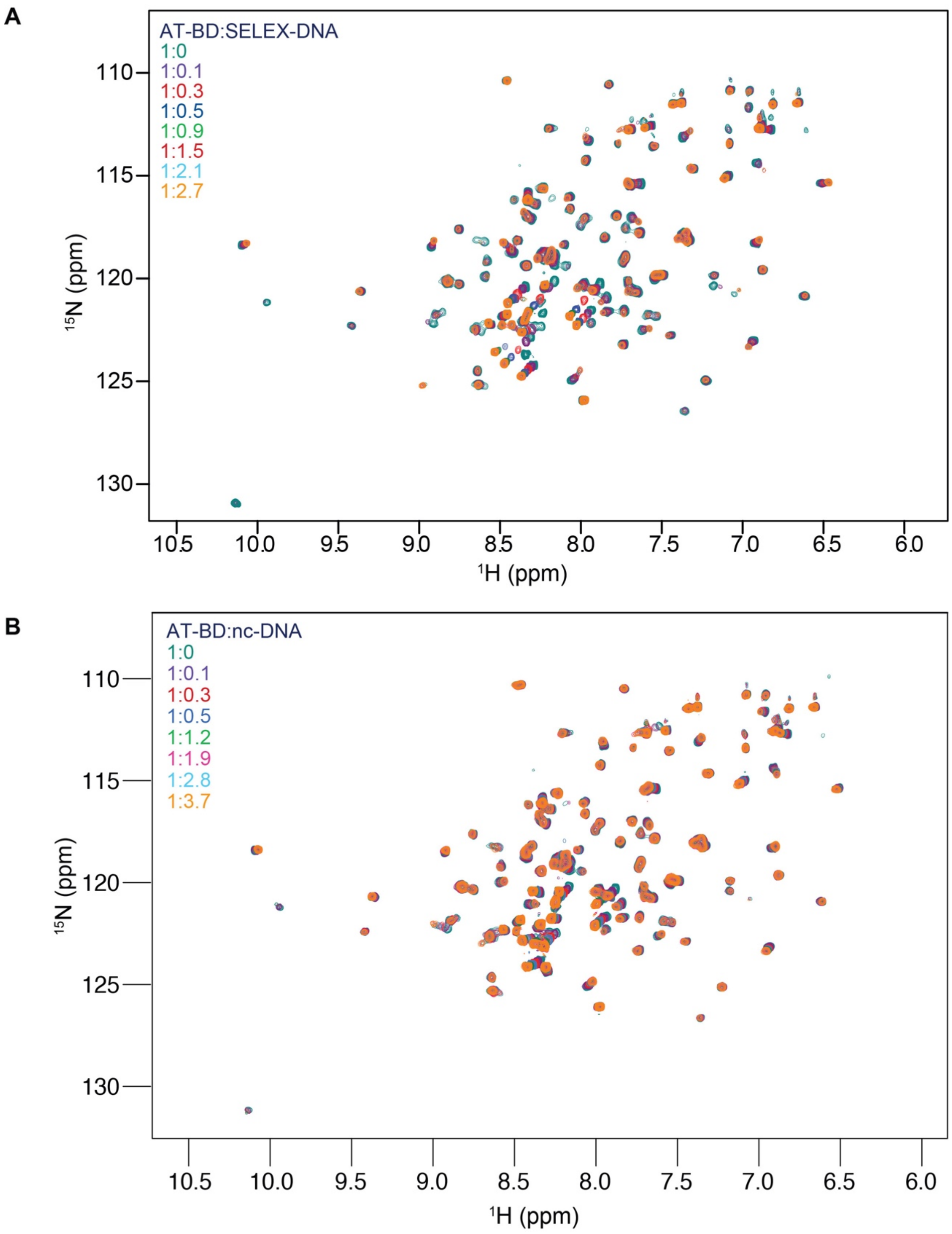


**Figure S2. NMR titrations with SELEX-DNA or nc-DNA. (A)** Overlay of $^{1}$H,$^{15}$N-HSQC spectra $^{15}$N-labeled AT-BD upon titration of unlabeled double stranded **(A)** SELEX-DNA or **(B)** nc-DNA.  and color coded according to the ratios indicated**.** Molar ratios are color coded according legend.

A

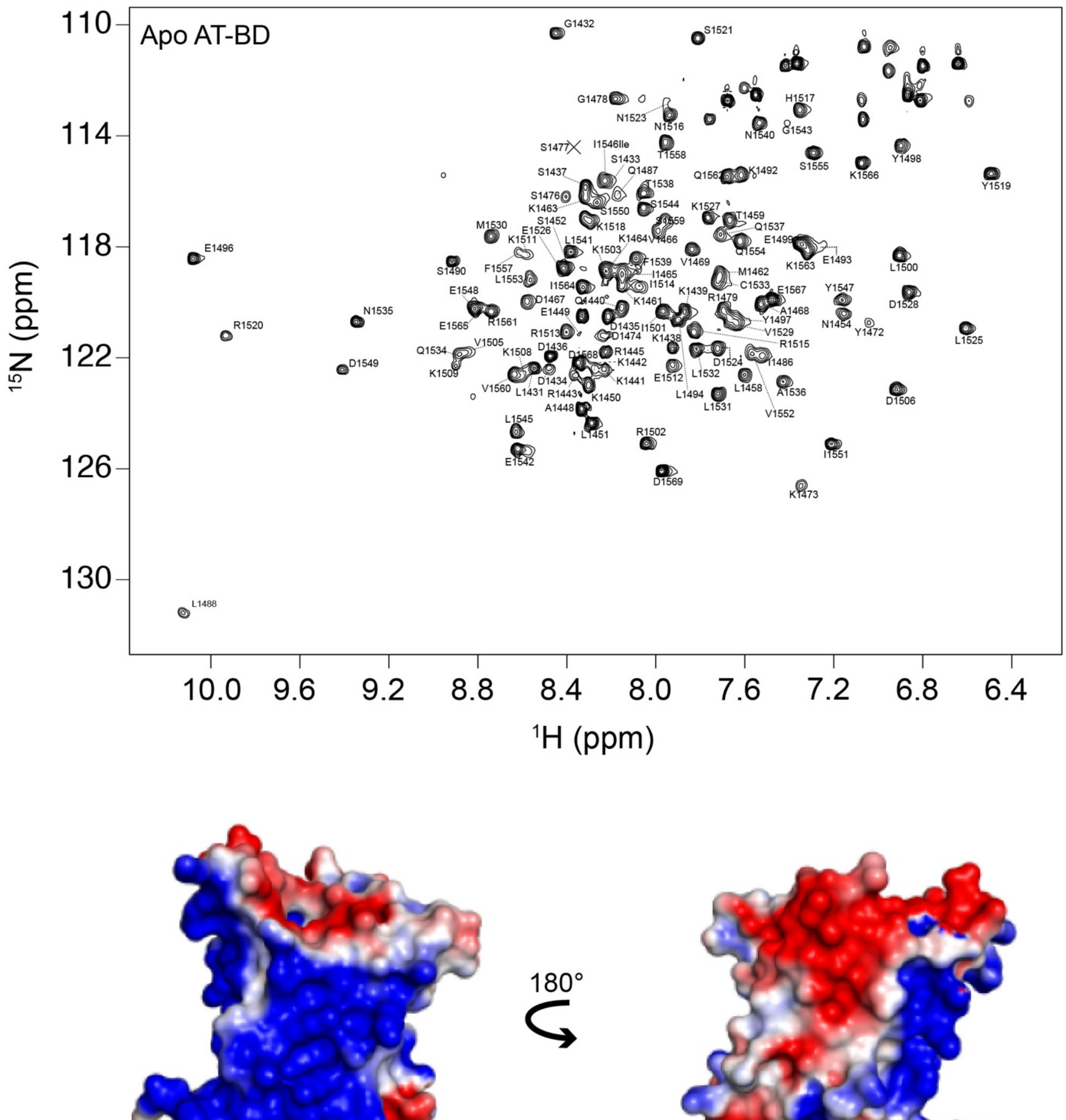


**Figure S3. Backbone assignment of the BRG1 ATBD. (A)** Final assignment for the backbone amide protons of AT-BD. The construct contains 141 residues, 9 of which are prolines. Thus, assuming fast exchange, 132 resonances would be expected. Of the 117 peaks observed 113 were successfully assigned. **(B)** Electrostatic surface potential for the BD (PDB ID 3UVD) as calculated by APBS in Pymol.

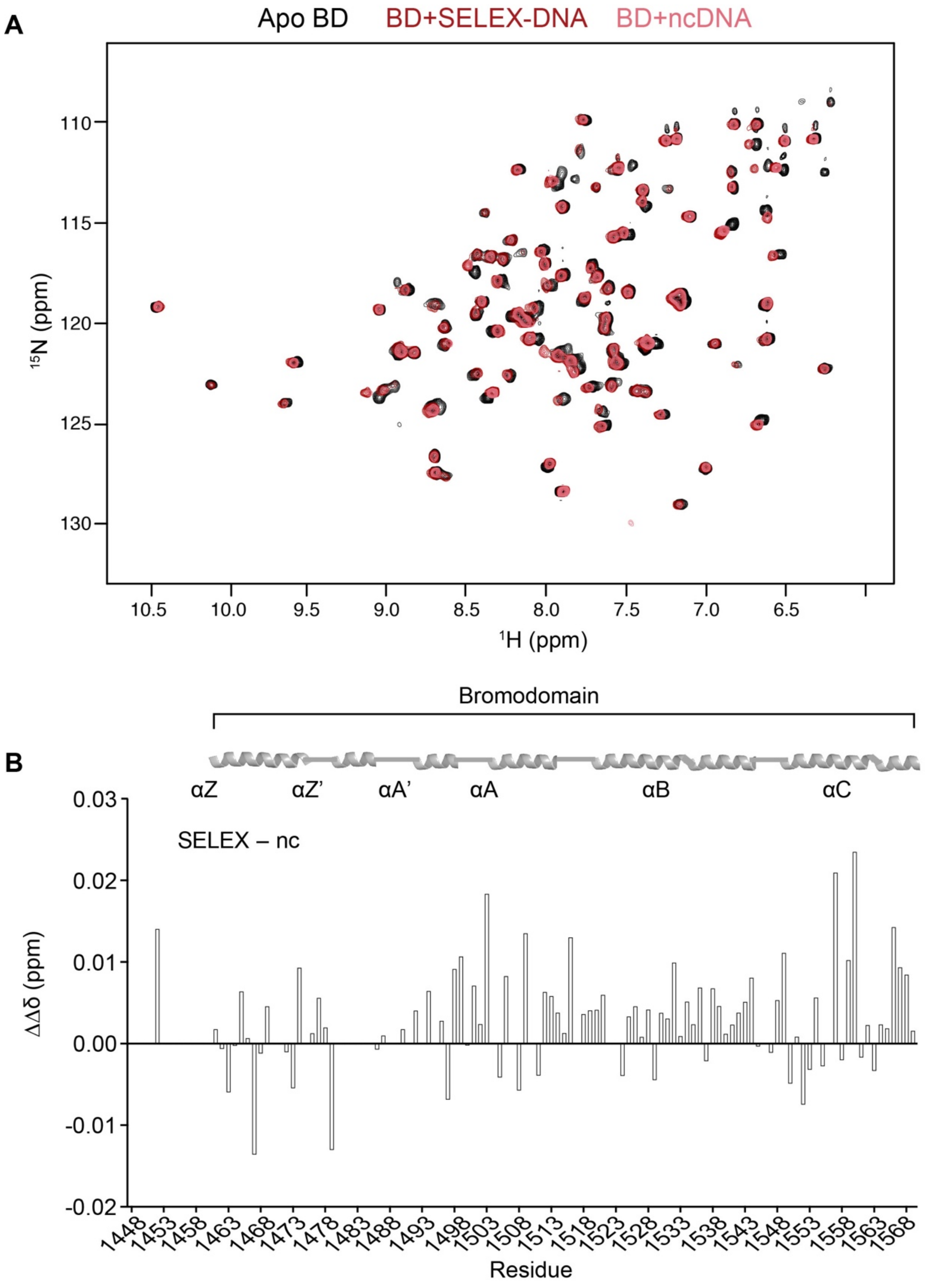


**Figure S4. The BD alone forms a more stable complex with SELEX-DNA. (A)** Overlay of $^{1}$H,$^{15}$N-HSQC spectra of $^{15}$N-BD (apo) saturated with SELEX-DNA (red) or nc-DNA (salmon). **(B)** Differences in the magnitude of CSPs (ΔΔδ) between bound states (SELEX-DNA minus nc-DNA) as a function of residue.

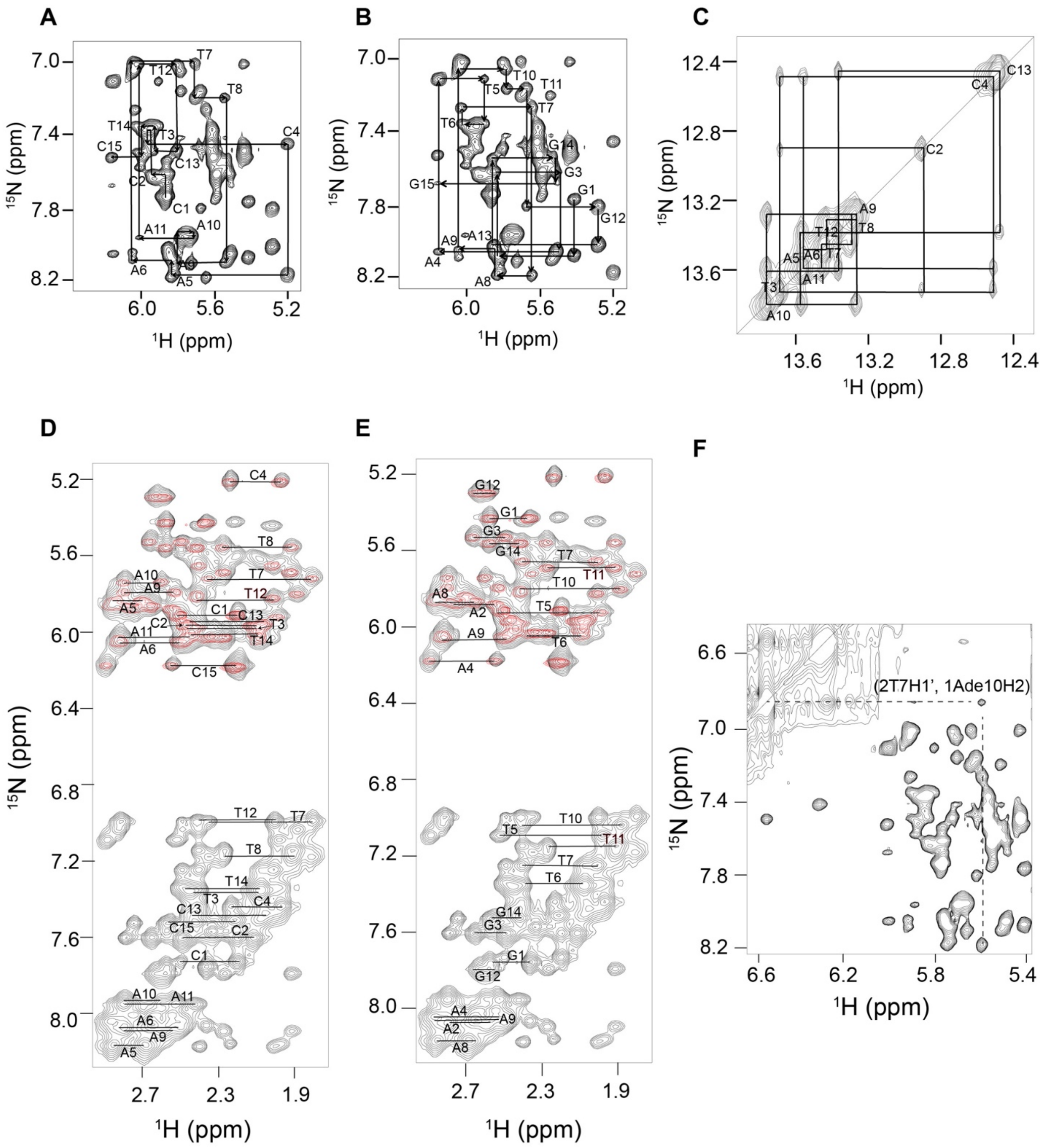


**Figure S5. Assignment of double-stranded SELEX-DNA. (A)** NOESY walk of the forward strand (H6,H8/H1′ area). **(B)** NOESY walk of the reverse strand. **(C)** Imino proton assignments. **(D)** Homonuclear NOESY spectrum (black) showing the assignments of the NOEs between H2′, H2′′ and H6/H8 protons (bottom) and NOEs between H2′, H2′′ and H1′ protons (top) for the forward strand and **(E)** for the reverse strand. TOCSY spectra (red) were also was used in assignments. **(F)** Inter-strand cross-peak NOE observed due to the narrowing of the minor groove in DNA.

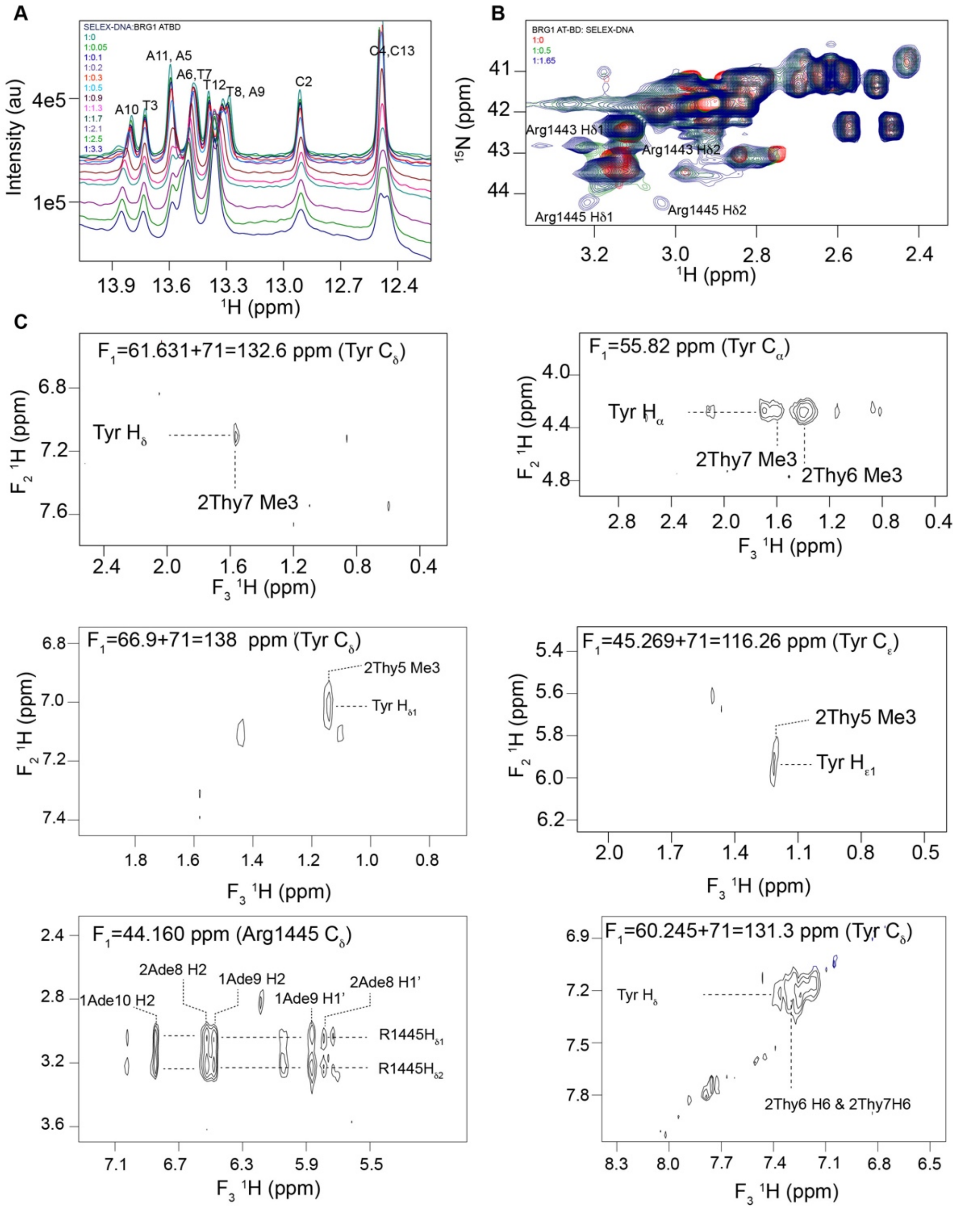


**Figure S6. AT-BD interaction with SELEX-DNA. (A)** Overlay of sequential 1D imino spectra of unlabeled double stranded SELEX-DNA upon titration with unlabeled AT-BD. Molar ratios are color coded as indicated in the legend. **(B)** Overlay of a selected region of $^{1}H$,$^{13}C$-HMQC spectra of $^{13}C$-AT-BD with unlabeled SELEX-DNA. Resonances for Arg1443 and Arg1445 split from single peaks in the apo state (red) to two peaks respectively upon saturation with SELEX-DNA (blue). **(C)** NOE crosspeaks between the AT-BD and DNA.

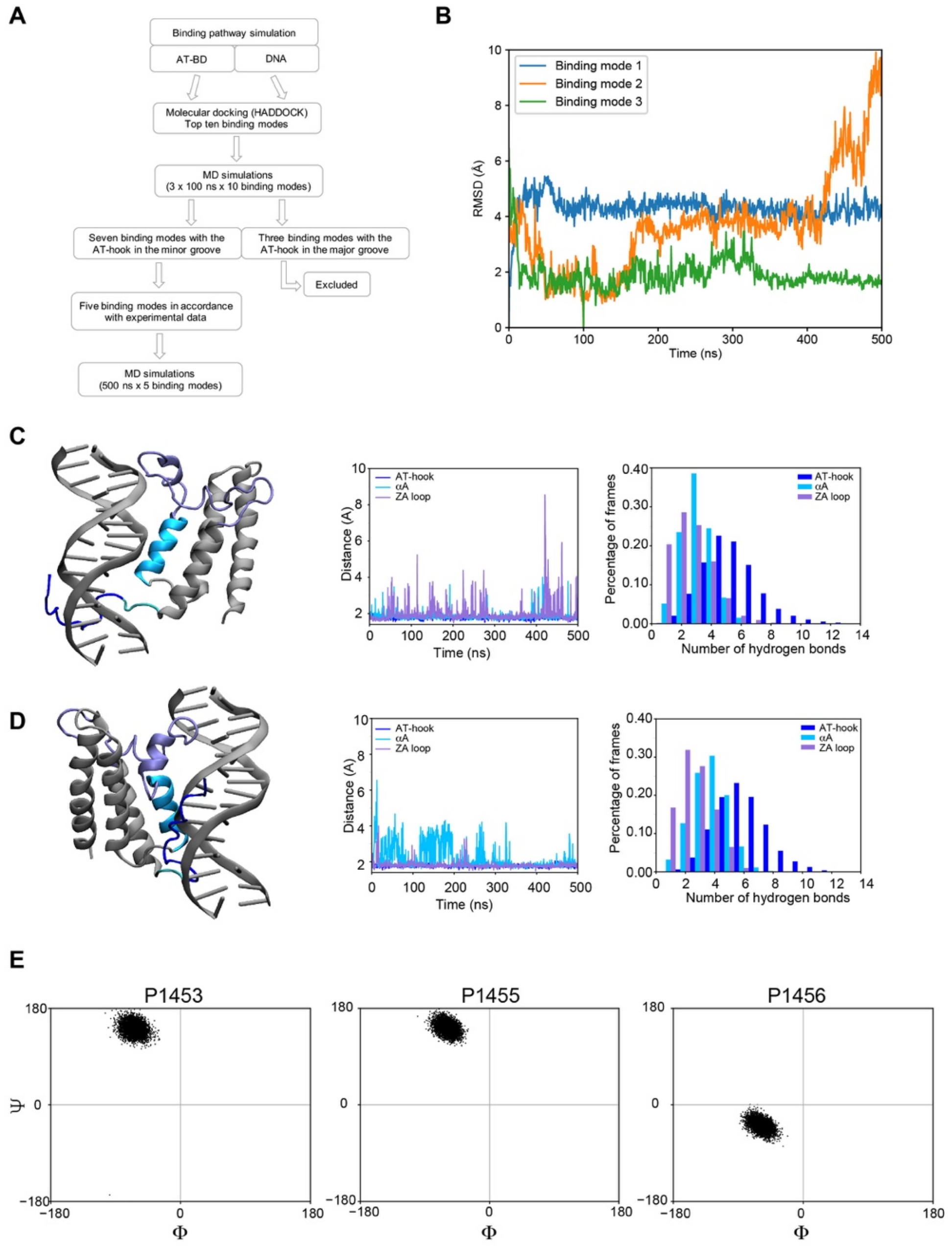


**Figure S7. Molecular model of the AT-BD/SELEX-DNA complex. (A)** Flow chart outlining the multi-step restraint-enabled refinement approach. **(B)** Root mean square deviation (RMSD) of AT-BD relative to SELEX-DNA for binding mode 1 (blue), 2 (orange), and 3 (green). **(C)** Model of AT-BD/SELEX-DNA for binding mode 2. The AT-hook is highlighted in blue, the αA in deep sky blue, the ZA loop in ice blue and the linker in cyan. The DNA and the rest of the BRG1 BD are in gray. The minimum distance between the AT-BD elements and DNA over the 500ns simulation are shown (middle). The number of hydrogen bonds between AT-BD elements and SELEX-DNA in the percentage of frames during the 500 ns simulations are shown (right). **(D)** The same as (C) for binding mode 3. **(D)** Computed Ramachandran angles for the three prolines in the linker (Pro1553, Pro1555, and Pro1556) over the course of the simulation.

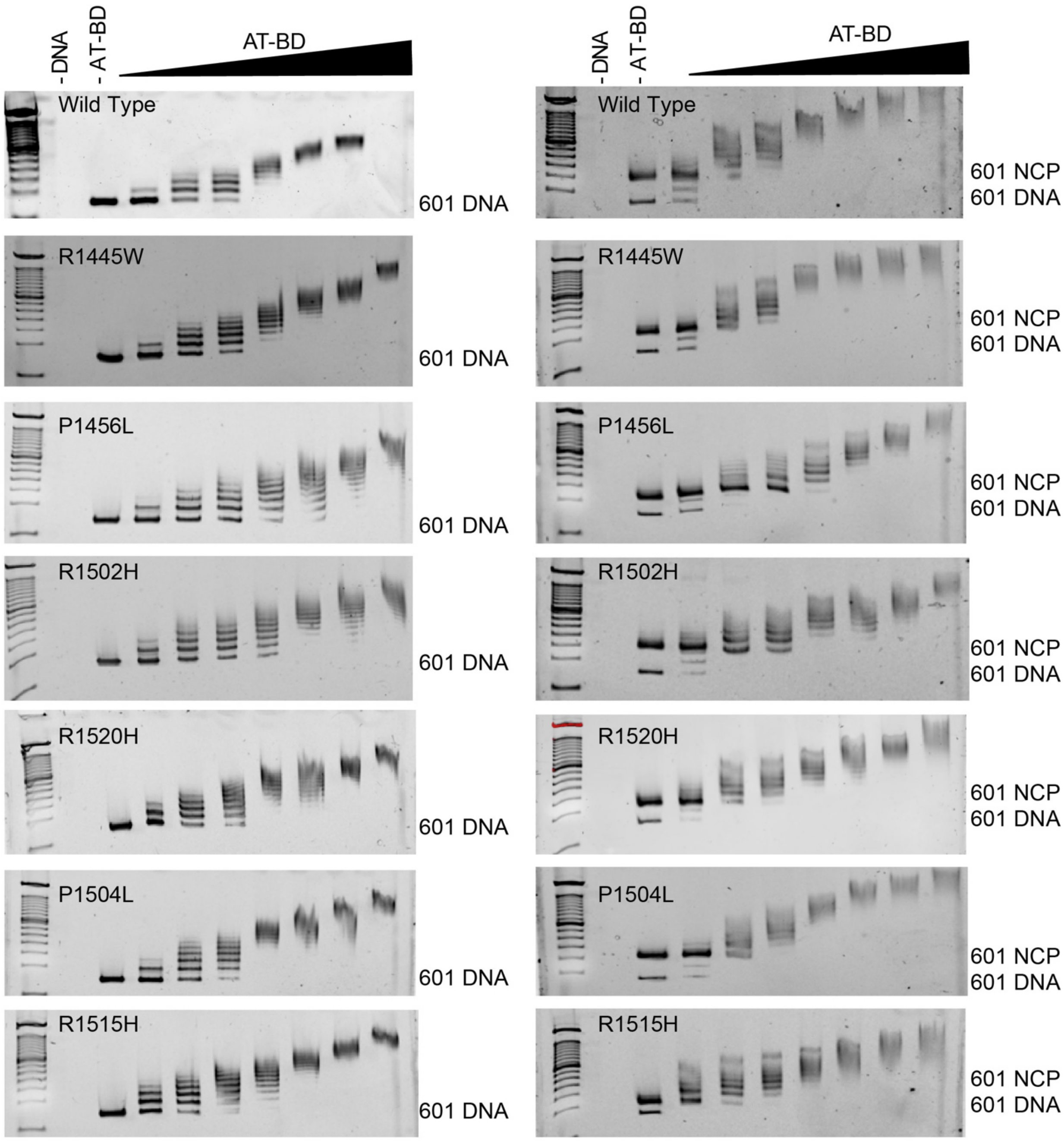


**Figure S8. Interaction of AT-BD cancer mutants with DNA and NCPs.** The interaction of wild-type (WT) AT-BD and AT-BD containing single cancer mutations were tested via EMSA. The 147bp Widom 601 DNA naked (left) or formed into nucleosome core particles (NCPs, right) was visualized on a native gel in the presence of increasing concentrations of WT AT-BD or AT-BD containing a single point mutation. Controls of protein with no DNA (-DNA) and DNA with no AT-BD (-AT-BD) were also run. DNA was visualized with Ethidium Bromide.

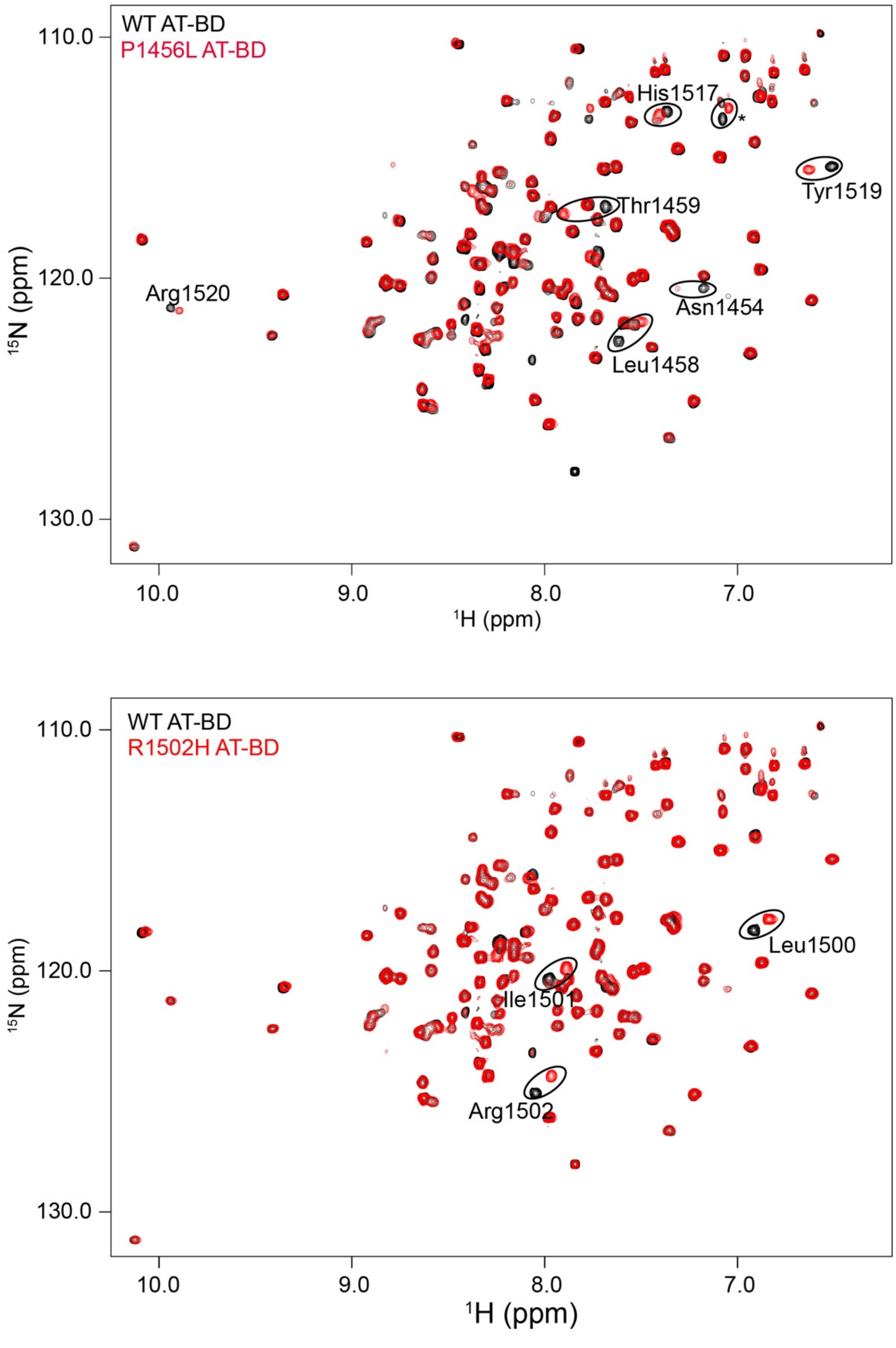


**Figure S9. Mutant protein folds.** Overlay of $^{1}$H,$^{15}$N-HSQC spectra of the apo states of WT AT-BD (black) and mutant (red) for P1456L (top) or R1502H (bottom). Residues that shift substantially are circled and labelled.
